# Trends in Subcutaneous Tumour Height and Impact on Measurement Accuracy

**DOI:** 10.1101/2022.09.29.510123

**Authors:** Daniel Brough, Hope Amos, Karl Turley, Jake Murkin

## Abstract

Tumour volume is typically calculated using only length and width measurements, using width as a proxy for height in a 1:1 ratio. When tracking tumour growth over time, important morphological information and measurement accuracy is lost by ignoring height, which we show is a unique variable. Lengths, widths, and heights of 9,522 subcutaneous tumours in mice were measured using 3D and thermal imaging. The average width:height ratio was found to be 1:3 proving that using width as a proxy for height overestimates tumour volume. Comparing volumes calculated with and without tumour height to the true volumes of excised tumours indeed showed that using the volume formula including height produced volumes 36X more accurate. Monitoring the width:height relationship (prominence) across tumour growth curves indicated that prominence varied, and that height could change independent of width. Twelve cell lines were investigated individually; the scale of tumour prominence was cell line-dependent with relatively less prominent tumours (MC38, BL2, LL/2) and more prominent tumours (RENCA, HCT116) detected. Prominence trends across the growth cycle were also dependent on cell line; prominence was correlated with tumour growth in some cell lines (4T1, CT26, LNCaP), but not others (MC38, TC-1, LL/2). When pooled, invasive cell lines produced tumours that were significantly less prominent at volumes >1200mm^3^ compared to non-invasive cell lines (P<0.001). Modelling was used to show the impact of the increased accuracy gained by including height in volume calculations on several efficacy study outcomes. Variations in accuracy contribute to experimental variation and irreproducibility of data, therefore we strongly advise researchers to measure height to improve accuracy in tumour studies.

## Introduction

Therapeutic efficacy of potential cancer therapies is commonly assessed by monitoring changes in subcutaneous tumour volume over time, by comparing tumour growth curves between control and treated groups of rodents. Accuracy is important both at small tumour volumes, when performing randomisation and determining the start of treatment and dosing, and at large volumes which are used when determining drug efficacy against controls, and to confirm if animal welfare points have been reached. Greater volume measurement accuracy can therefore give greater confidence in results with the potential to run shorter studies, or requiring fewer repeats to achieve the same level of statistical significance in results.

Important information about the tumour is lost when recording only tumour length and width over time, as is standard in preclinical studies. Subcutaneous tumours are inoculated below the dermis and grow outwards, protruding from the rodent body. On average, the tumour shape can be generalised as an ellipsoid where width and height are equal(1), however the shape and prominence (width:height relationship) of subcutaneous tumours can be affected by several variables including injection technique, cell line, tumour type, and growth stage. Large subcutaneous tumours for example, are known to have a non-spherical, flattened shape(2). Complex tumours may have multiple lobes (3), and tumours where a necrotic core develops may also collapse in on themselves, producing a ‘donut’ or ‘cratered’ shape (4). Tumours with an aggressive, metastatic phenotype gain the ability to invade into other tissues during the transition to metastasis (5). We might reasonably expect that this local invasiveness at the primary site would be detectable by tracking tumour prominence over time.

Tumour type and cell line also affects subcutaneous tumour shape. Lymphoma tumour cell lines form tumours with less stroma than other tumour types including carcinomas and produce characteristic ‘diffuse’ or ‘flattened’ subcutaneous tumours that can be distinguished from carcinomas by eye(6). Anecdotally, technicians have reported such diffuse, flat tumours are difficult to measure consistently using standard calliper techniques due to their poorly defined boundaries that are difficult to define by eye and surround with calliper blades.

The low-cost, high throughput method of choice for capturing tumour volume is manual measurement with callipers. The longest tumour length and width measurements (as judged by the user) are taken by placing calliper blades on the outside of the tumour bulk protruding from the mouse body. Calliper measurement is known to introduce bias as the direction to measure the longest length is subjective, especially in tumours with complex morphologies (5).Calliper contact with the tumour can also cause squeezing of the tumour which leads to underestimation of the true volume.

Height is difficult to capture using callipers as the tumour base is obstructed by the mouse body, leading to larger coefficient of variation and instrumental error from height measurement(2). Measurement of height also increases animal handling time when using callipers. Thus, methods to estimate tumour volume from length and width alone are used most commonly, including the LWW formula which uses tumour width as a proxy for height in a 1:1 ratio to estimate tumour volume (8,9). This formula does cause overestimation of tumour volume however as it does not take into account any changes in height which is smaller than width(5). Different approaches have been suggested in order to achieve more accurate tumour volumes without measuring height(2,9) but these methods have not been widely adopted or implemented.

### LWW Formula

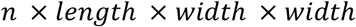

where *n* is a constant, most commonly π/6 (used here) or 0.5.

Alternative measurement devices offer increased accuracy and reduced bias including CT-(7), fluorescence-(10), Magnetic Resonance-(11), and ultrasound imaging(12). These techniques allow visualisation and measurement of the whole tumour in situ, however the main disadvantages; cost, low throughput, and significant welfare risks from anaesthesia and invasive techniques ensure that calliper measurement remains the standard for subcutaneous tumour measurement.

3D and thermal imaging (3D-TI) is a technique that captures and records tumour morphology for volume calculation(13). In comparison to callipers 3D-TI reduces user bias by using machine learning and computational algorithms to calculate length, width, and height measurements automatically. The variability in calliper tumour measurement methods is predicted to affect the rate of false negative results in efficacy studies ((14), paper in review), and the tumour growth inhibition (TGI) drug efficacy metric(15) in comparison to 3D-TI. Height measurement allows 3D-TI to calculate tumour volume using the LWH formula which has been shown to give more consistently accurate volumes in comparison to the LWW formula and 18 other tumour volume formulae(8). The LHW formula has also been proven to improve accuracy in clinical studies(16).

### LWH Formula(8)

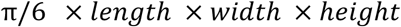

### Aims

Including tumour height in volume calculations has been shown to produce more accurate tumour volumes than formulae using length and width dimensions alone. We aimed to use our anonymised global database of tumour scans taken by 3D-TI users to first define the average relationship between tumour height and width, then aimed to validate our findings by comparing LWH and LWW tumour volumes to the true volumes of excised tumours.

The second part of the investigation aimed to investigate the real-world impact of variations in tumour prominence in more detail, by breaking cell lines down into metastatic vs non-metastatic groups, and grouping by tumour types. To quantify the importance of measuring tumour height during studies, in silico modelling was used to simulate the effect on efficacy study outcome metrics (TGI, false positive rate, and false negative rate) when comparing LWW and LWH measurement formulae.

## Methods

### In vivo data collection

The 3D-TI device and system (BioVolume) and support were provided to 27 client organisations by BioVolume Ltd (previously Fuel3D). Subcutaneous tumour models were established in the mouse flank area from inoculated cell lines according to client organisation protocols. Training was provided by BioVolume Ltd over a series of sessions either in person or online. Mice were scruffed according to client animal handling protocols, and the tumour region was presented to the BioVolume silicon aperture for image capture (0.5s). Tumour length, width, and height were measured automatically by the 3D-TI device as previously described(13).

Methods and welfare endpoints adhered to US or EU welfare and ethics rules depending on lab location. All animal care, lab work, calliper measurements, and 3D-TI scans were carried out by scientists in client organisations. Data were shared with BioVolume (previously Fuel3D) to use in an aggregated and anonymised way. Client companies and scientists did not have financial interests in BioVolume.

Maximum tumour volume limits are typically 1.2-1.5cm diameter in the UK(17) and 2cm diameter or ~2000mm^3^ in the US, therefore we looked at measurement accuracy up to ~2000mm^3^.

### Data Processing

#### Evaluation Dataset

Our global dataset of tumour scans included 39,758 scans from 119 cell lines and 156 users in 27 organisations. These were filtered to include data from evaluation studies only, as these studies were performed after user training was completed, and a high level of scan quality was maintained by users. This comprised 9,522 scans from 16 cell lines and 46 users (Evaluation Dataset). Only established cell lines with no additional genetic modification were included. To better account for any non-linearity, a generalised additive model (GAM)(18) was applied to this dataset to better understand the relationship between height and width across different tumour volumes.

#### Control Dataset

Scans in the Global Dataset were assigned to a Study and Group by users during scanning. We identified control groups in the Global Dataset using group names. Scans in other groups were filtered out, to reduce potential effects of treatments on tumour morphology and to enable like-for-like comparison between untreated tumour cell lines. Any ambiguously labelled groups were excluded and only immortalised cell lines were included. 42 cell lines and 9,067 scans were present in this dataset (Control Dataset).

#### Excised Weights Dataset

The global dataset was filtered to remove any scans where excised weight was not available. This dataset consisted of 299 scans and 274 corresponding calliper measurements across 13 studies, 12 cell lines, and 6 mouse strains.

#### Modelling

Testing confirmed that the residuals of all models were normally distributed, thereby meeting the assumptions of linear modelling. To verify that models described the data well, predicted tumour measurements were plotted against the actual tumour measurements for both models, a y=x line was plotted and used to assess model fit (all available in Supplementary Materials).

### Tumour Prominence by Cell Line

To begin the investigation into tumour height, an initial exploratory plot using the Evaluation Dataset scans was created to observe the width:height (prominence) relationship across all 16 cell lines. Tumour height was plotted against tumour width as measured by 3D-TI. A generalised additive model with integrated smoothness integration using the “REML” method (19,20) was then fitted to all the Evaluation Data using the ggplot2 geom_smooth function in R(19) with an intercept set through the origin to see if there was a linear relationship between the height and width, along with a 95% confidence interval associated with the prediction of the linear model.

Six immortalised cell lines in the Evaluation Dataset were chosen to study in greater detail: 4T1, CT26, LNCaP, MC38, RENCA and TC-1 (Table 1). They were amongst the most frequently measured tumour cell lines, and all represent carcinoma tumours. A linear model was fit (using stat_smooth (19)) to each cell line to assess width:height relationship, along with a 95% confidence interval associated with the prediction of the linear model.

**Table 1.**
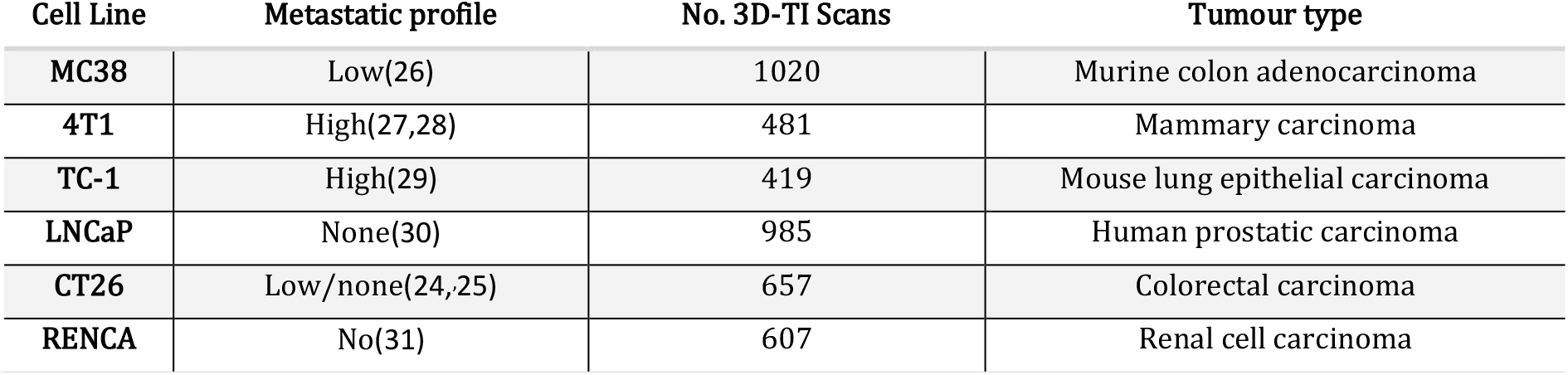
Metastatic profiles of subcutaneously inoculated tumour cell lines used when comparing cell lines. Numbers in brackets refer to bibliographic references.

| Cell Line | Metastatic profile | No. 3D-TI Scans | Tumour type |
| --- | --- | --- | --- |
| MC38 | Low(26) | 1020 | Murine colon adenocarcinoma |
| 4T1 | High(27,28) | 481 | Mammary carcinoma |
| TC-1 | High(29) | 419 | Mouse lung epithelial carcinoma |
| LNCaP | None(30) | 985 | Human prostatic carcinoma |
| CT26 | Low/none(24,25) | 657 | Colorectal carcinoma |
| RENCA | No(31) | 607 | Renal cell carcinoma |

Following the initial exploratory plot and the detection of a non-linear height:width relationship, volume bins were used to group tumours by size for a new linear model. The lm function from the stats package in R was used (21). This way prominence was directly compared across cell lines without the influence of tumour size, providing more confidence in the results that any differences would be down to the cell line. No intercept term was used and the global intercept was set to 0 since a tumour of 0 width should also have 0 height, which in turn meant that the fit would have to pass through the origin. Data were removed for cell lines and bins combinations that contained less than 3 datapoints as this was not enough to create a linear model.

The model formula is outlined below:

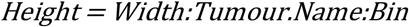

Control group data was used in order to remove any effects that treatment may have had on prominence. The new linear model was then applied to the same immortalised cell lines (4T1, CT26, LNCaP, MC38, RENCA and TC-1) as previously mentioned, with the number of scans for each cell line outlined in Table 1. 95% confidence intervals associated with the slope estimates were also provided. A two-way ANOVA (22,23) was computed for unbalanced designs (type III) to test whether the variability between cell lines was greater than the variability within cell lines.

### Excised Weights

To calculate tumour weights, tumours were excised on the final measurement day. The true volume of excised tumours was derived by weighing the tumour, then volume was calculated in a 1:1 ratio (mg to mm^3^). The difference between calculated volume and excised weight was calculated in a relative fashion (percentage difference) to compare the differences between the volumes obtained.

Using the excised weights dataset, prominence (height divided by width) was calculated from the 3D-TI measurement and matched with the corresponding calliper measurement where data were available, then plotted against the percentage difference from excised weight. A linear model was then fitted to the graph along with the error associated with the model.

### Prominence by Metastatic Profile

Cell lines studied previously were classified by metastatic profile according to reports in the literature (Table 1). Only metastasis from tumours inoculated subcutaneously in mice was considered to be relevant to our study; metastasis from intravenous or orthotopic inoculation was discounted. Cell lines where a definitive classifier could not be found in the literature were excluded. CT26 was both reported as non-metastatic(24), and to have some incidences of metastasis from subcutaneous tumours <2000mm^3^ which increased over time(25). MC38 was characterised as having a low rate of metastasis from subcutaneous sites because metastasis rate was <50% although final tumour volumes were much larger than today’s permitted maximum volume endpoints(26).

3D-TI data from the Control Dataset were sorted into ‘Metastatic’ and ‘Non-Metastatic’ groups in the same way. Data were filtered such that any cell lines with uncertainty surrounding their metastatic nature were excluded e.g., conflicting results in the literature. Cell lines with <150 tumour scans were also excluded, leaving 5 metastatic cell lines (1771 scans) and 5 non-metastatic cell lines (2380 scans) (Table 2). Data from this filtered dataset were plotted and two linear models were fitted, with 95% confidence intervals also being provided.

**Table 2.** Cell lines used to compare prominence trends between metastatic and non-metastatic cell lines. References are included. Numbers in brackets refer to bibliographic references.

| Tumour Name | Metastasis from Subcutaneous Site | Scans |
| --- | --- | --- |
| HCT116 | No (33) | 188 |
| LNCaP | No (30) | 985 |
| MCF7 | No (34) | 228 |
| MiaPaCa2 | No (35) | 376 |
| RENCA | No (31) | 607 |
| 4T1 | Yes (27,28) | 481 |
| B16F10 | Yes (29) | 465 |
| H358 | Yes (36) | 243 |
| LL/2 | Yes (37) | 176 |
| TC-1 | Yes (29) | 419 |

The model formulae are outlined below:

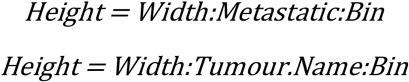

This was followed by computation of a two-way ANOVA (23) for unbalanced designs (type III) and a Tukey’s honest significant differences test. A general linear hypothesis test in the Multcomp R package (32) was then also conducted to test for significant differences in slope estimates of the model between the two groups for each of the individual volume bins.

### Prominence by Tumour Type

To investigate whether cell lines of the same tumour type displayed similar prominence characteristics, cell lines within different groups from the Control Dataset were compared - these subgroups were composed of carcinomas (27), gliomas (1), melanomas (4), leukaemias (3), and lymphomas (2). In order to give a fair representation of each cell type, cell lines with less than 9 datapoints in total were excluded. The control group data for carcinomas vs non-carcinomas were fed into a height:width linear model accounting for carcinoma status (yes or no) and each volume bin.

The model formula is outlined below:

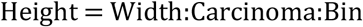

A two-way ANOVA (23) for unbalanced designs (type III) and a Tukey’s honest significant differences test were computed. A general linear hypothesis test in the Multcomp R package (32) was then used to test for significant differences between each of the volume bins.

### Generating the In Silico Efficacy Dataset

To outline the impact that failure to measure height could have on an efficacy result, the width-height models using volume bins described in the ‘Tumour Prominence by Cell Line’ section were used to create an artificial dataset for 4T1, CT26, LNCaP, MC38, RENCA and TC-1. Synthetic data were generated with the aim of removing any unwanted sources of variability that come with the aggregated data such as measurements coming from different studies and laboratories and to also generate a larger sample size. To consider cell line-specific growth rates, real data were fed into separate linear models for the length and width tumour dimensions, accounting for change in growth on a daily basis and creating an intercept for each study since not all studies started at the same volume (only studies with greater than 3 measurement sessions were used).

The model formulae are outlined below:

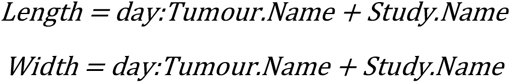

The obtained growth rates were then multiplied by a list containing the unique days within the synthetic study (multiples of 3 starting from 3 through to 90). Volumes were calculated using the LWW formula and a maximum volume limit of 2500 mm^3^ was set. From these data, 10 synthetic rodents were generated and initial tumour volumes were offset by applying the same standard deviation to length and width as observed within the control group data for days and studies where the LWW volume was less than 120mm^3^.

The individual rodent growth rates for length and width (gradients) were varied by a standard deviation equal to the slope error observed when calculating the growth rate coefficients for each cell line. The LWW volume for each rodent was then recalculated and results were generated. Lengths and widths that were less than 0 were excluded from the dataset. The mean for each cell line on each day was calculated at this point. The data were then duplicated to create a treated group with identical starting conditions, with ‘treatment’ applied on the first day after the group mean exceeded 80mm^3^. ‘Treatment’ was simulated by multiplying individual length and width dimensions of the treated group tumours by a defined treatment strength (from 0.5 to 0.9), with 0.5 being the strongest treatment level and 0.9 being the weakest. This had the effect of reducing the length and width growth rates by 100*(1 – treatment strength) %, for example; at a treatment level of 0.9 the growth rate would be reduced by 100*(1 – 0.9) = 10% for the treated group beginning on the measurement session after first treatment was applied. Within this step, tumour heights were predicted by fitting data to the linear model for the cell lines and the LWH volume was computed. This step was repeated across 41 varying levels of treatment strength. This process was then repeated 250 times after generating 250 sets of 20 animals (10 control, 10 treated).

### Using the In Silico Dataset to Calculate Tumour Growth Inhibition

The ratio of final volume to treated day volume was calculated for each rodent for LWW and LWH volumes. For each iteration, cell line, volume type, and efficacy level the tumour growth inhibition (TGI) was computed by dividing the mean rodent ratio for the treated group by that of the control group (group sizes were both equal). To show this graphically, the mean TGI was computed across the iterations and plotted with 95% confidence intervals against the efficacy level for each volume type and cell line. To calculate the number of successful results, a conservative TGI threshold of 0.45 was used and applied such that any result under 0.45 would be considered to have shown effectiveness of treatment and any result above 0.45 would show ineffective treatment. The LWH volume was taken to be the ground truth in this scenario as it was established to be the most accurate volume formula. The number of successful trials were calculated using the TGI cut-off (<0.45) and the results for each volume type were compared to each other. LWW error rates were calculated by comparing the number of successful trials in comparison to LWH for each cell line. For example, if LWW had a greater number of successful trials than the LWH standard then this would indicate false positive results in some LWW trials.

## Results

### The Tumour Height:Width Relationship

Plotting tumour height against width clearly showed that width is not a good proxy for tumour height, with none of the 9,522 tumour height values being equal to or exceeding their corresponding widths (Figure 1A). Height was roughly 1/3 of the width on average: Height(mean) = (0.325 ± 0.001)*Width(mean). The heteroskedastic nature of the plot means that as width increases, so do the range of possible height values for a given width. This reveals the unique and independent behaviour of tumour height, further outlining why using width as a proxy for height is not an accurate approach when calculating tumour volume. The next step was to focus on the height:width relationship for individual cell lines.

**Figure 1.**
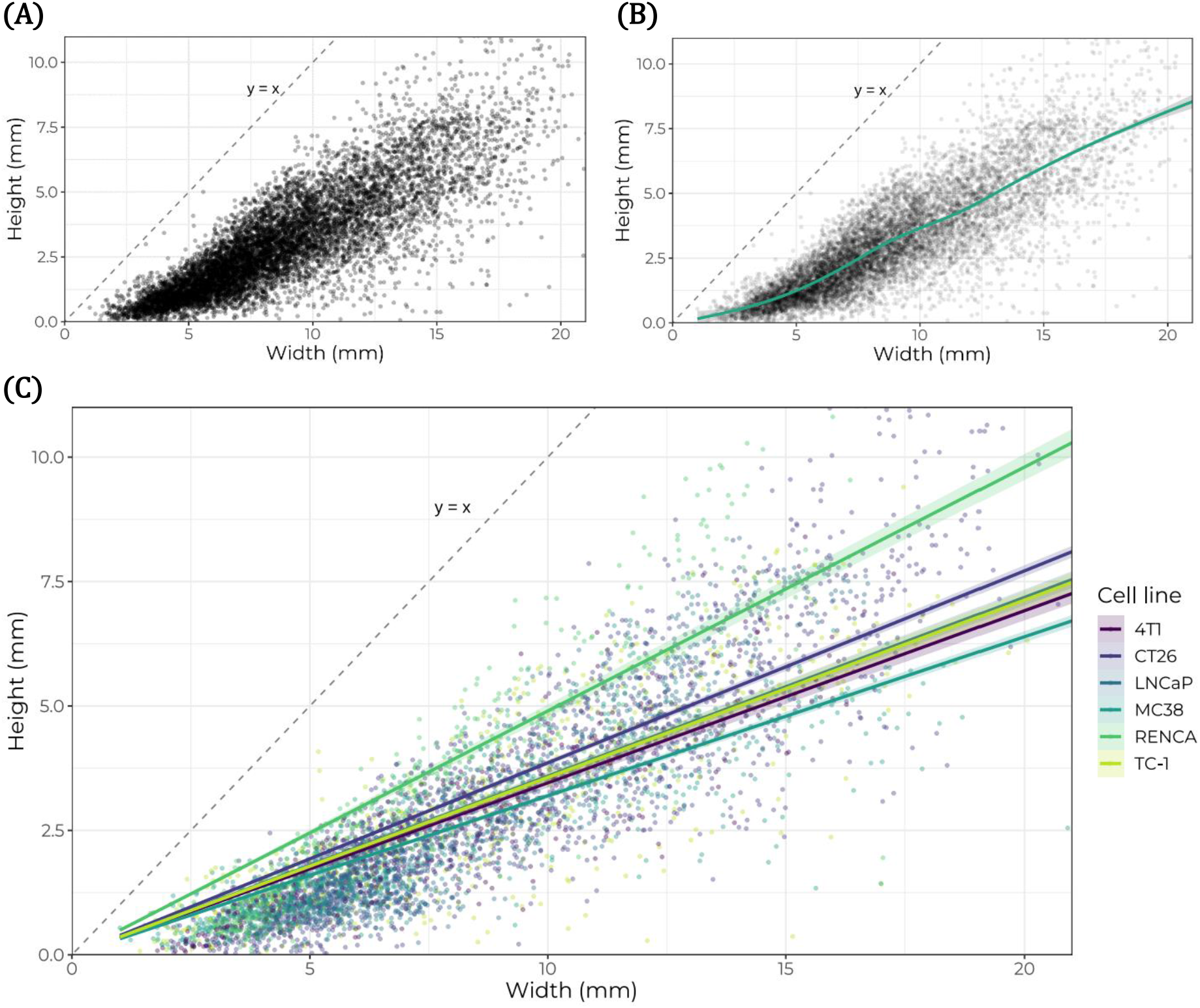
Investigation of subcutaneous tumour prominence. (A)Height vs width, n=9,522 3D-TI scans (Evaluation Dataset). Axes capped at (11mm and 21mm). (B)Generalised additive model with integrated smoothness integration using the “REML” method(19,20) (green line) plotted on Figure 1a data, along with a 95% confidence interval associated with the prediction of the linear model (shaded). (C)Height vs width for subcutaneous tumours of 6 cell lines (4T1, CT26, LNCaP, MC38, RENCA, TC-1. N=5,342 total). Trend lines were fitted using a linear model for each cell line. Shaded line region shows the 95% confidence level associated with the prediction of the linear model. Y=x shown as dashed line on each plot for reference.

### Tumour Prominence by Cell Line

Data for six cell lines (4T1, CT26, LNCaP, MC38, RENCA and TC-1) were plotted to investigate any unique cell line characteristics (Figure 1C). The tumours from different cell lines varied in prominence with respect to each other; RENCA had the greatest prominence (height/width) of the six tumours, with MC38 forming the least prominent (flattest) tumours. The linear model appeared to underfit by a large margin at smaller widths, suggesting a non-linear distribution of the data.

To confirm this hypothesis, a generalised additive model was fitted to the evaluation dataset (shown in Figure 1A) which, following observation of the fitted GAM slope, suggested a non-linear height:width relationship at smaller volumes (Figure 1B). This indicates that on average tumours were flattest (least prominent) at the earliest stages of growth, below widths of 8mm.

### Prominence Trends Across Increasing Tumour Volumes

A new model introducing volume bins was applied to individual cell line data to determine if this non-linear height:width relationship was cell line-specific (Figure 2). The different trends for each of the cell lines show that the height:width ratio (prominence) can change as a tumour grows and that cell lines exhibit their own unique distributions. 4T1, CT26, LNCaP, and RENCA tumour volume increase correlated with an increase in prominence up to ~1200mm^3^, however MC38 and TC-1 started off as relatively more prominent tumours but the height:width relationship then stayed relatively stable over the same range. RENCA tumours exhibited the fastest increase in height relative to width up to a peak at 1200-1600mm^3^. This varying ratio of height:width within a given cell line could impact efficacy results, as key height differences between groups could go unnoticed when only measuring length and width.

**Figure 2.**
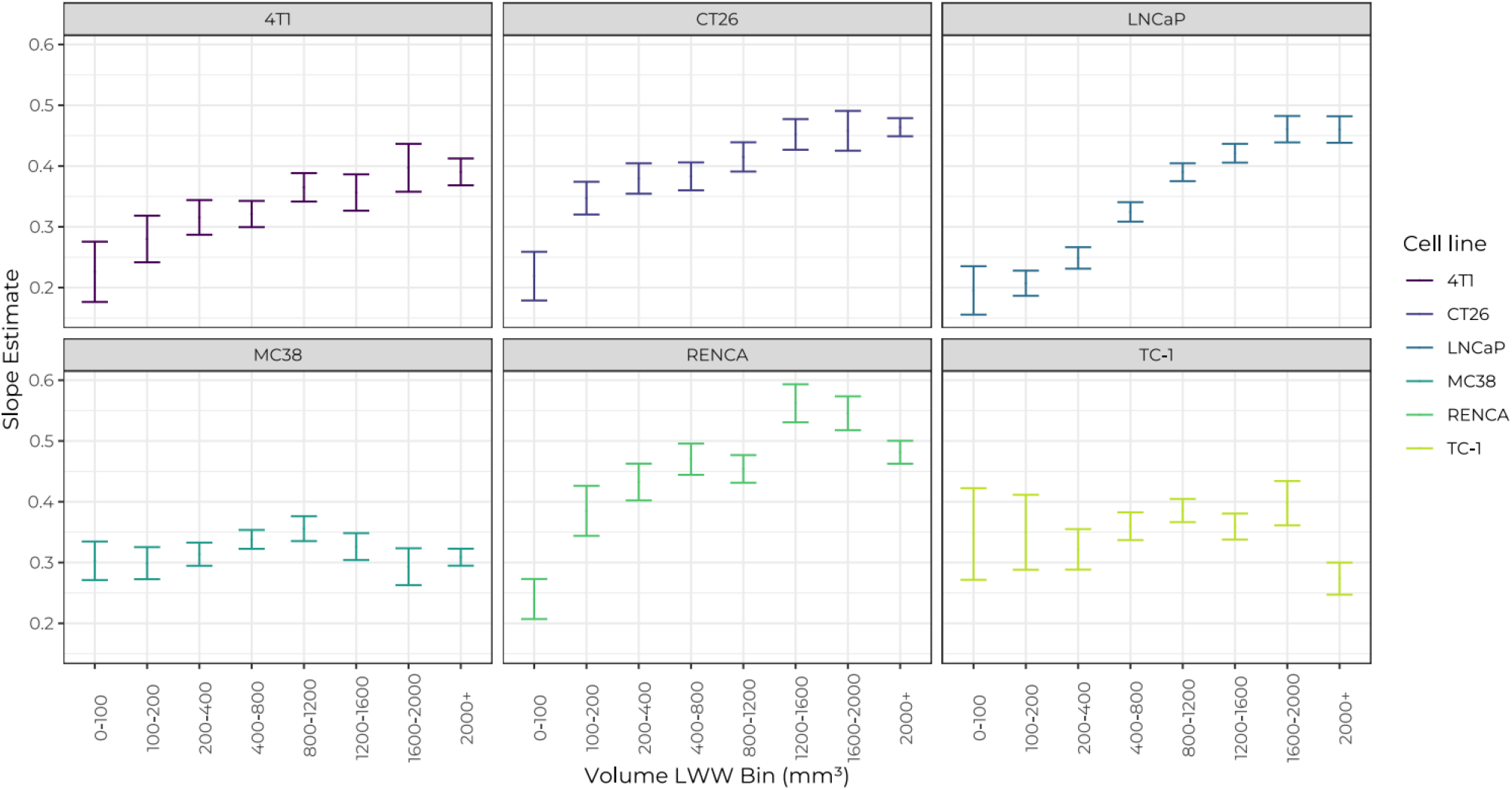
Tumour prominence by cell line. Slope estimates of the Height = Width:Tumour.Name:Bin model for cell lines 4T1, CT26, LNCaP, MC38, RENCA, and TC-1. Error bars represent the 95% confidence intervals of the slope estimates.

The individual distributions of tumour prominence by cell line and bin size were plotted to compare differences between cell lines (Figure 3). Two-way ANOVA yielded a p-value of P<0.0001 across the cell lines, volume bins and their interactions, showing that the variability across cell lines was significantly greater than the variability within them. Cell line pairs were also compared by ANOVA (Table 3) with statistical difference detected between cell lines such as CT26-4T1, but not for other pairings such as LNCaP-4T1. This indicated that tumour prominence is dependent on tumour cell line.

**Table 3.** Comparison of cell line tumour prominence. Tukey honest significant differences test computed using fitted ANOVA to give multiple pairwise-comparison between cell line group means. Significance indicated: P>0.05 ns, P≤0.05 *, P≤0.005 **, P≤0.001 ***

|  | 4T1 | CT26 | LNCaP | MC38 | RENCA | TC-1 |
| --- | --- | --- | --- | --- | --- | --- |
| 4T1 | x | *** | ns | ns | *** | *** |
| CT26 | *** | x | *** | *** | ** | ns |
| LNCaP | ns | *** | x | *** | *** | *** |
| MC38 | ns | *** | *** | x | *** | *** |
| RENCA | *** | ** | *** | *** | x | ns |
| TC-1 | *** | ns | *** | *** | ns | x |

**Figure 3.**
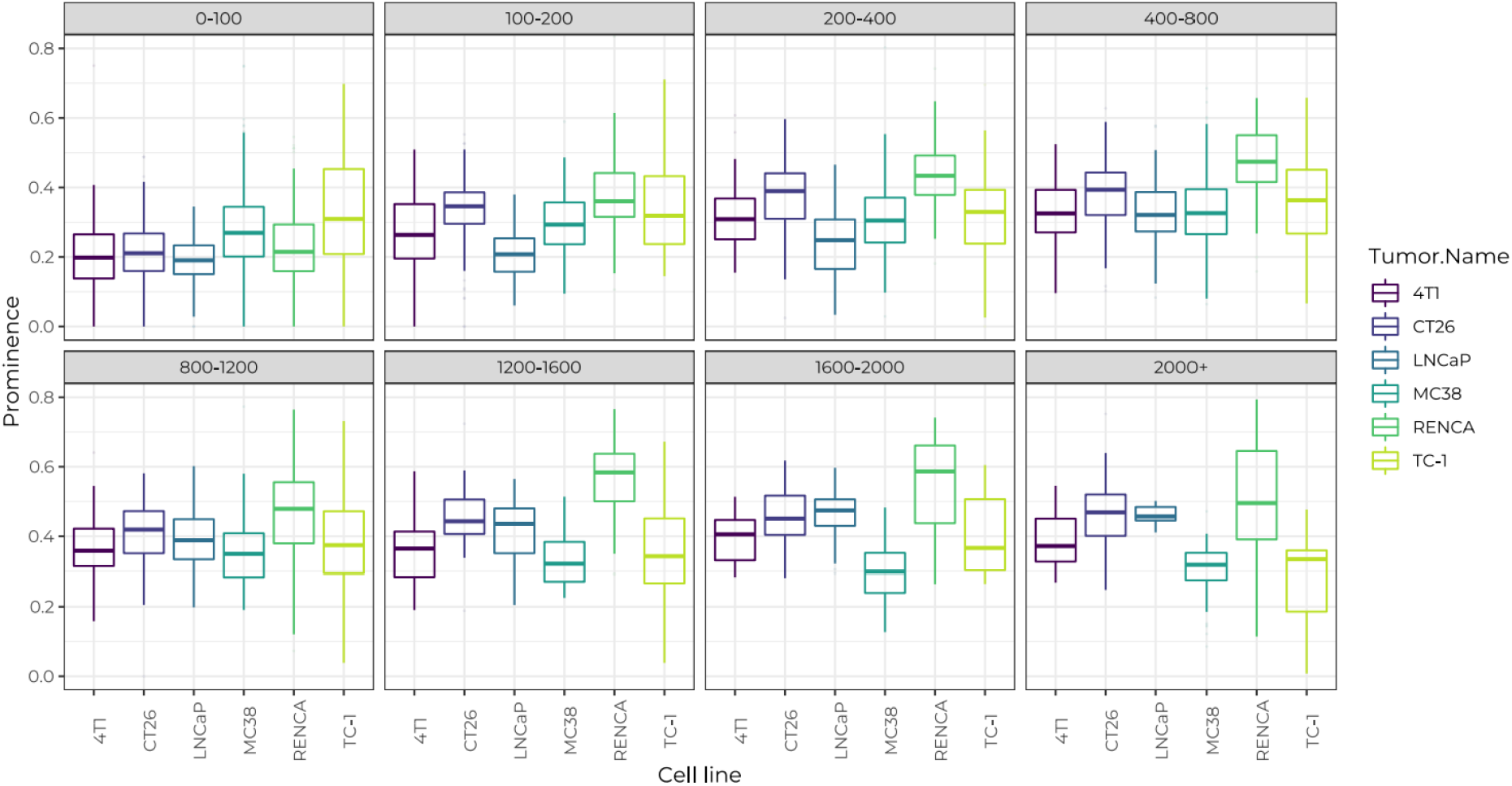
Prominence value distributions for each cell line. Distributions are shown as box plots separated out by LWW volume bin (mm^3^).

4T1, LNCaP and MC38 tumours were not significantly different to each other. TC-1 and CT26 were not significantly different to each other. RENCA tumours were significantly different to all other cell lines which was unsurprising as they had a higher median prominence than all other cell lines above 100mm^3^, indicating that RENCA tumours had the most prominent morphology (Figure 5).

### Investigating Metastatic Cell Lines

We hypothesised that metastatic tumours could affect accuracy of measurements due to aggressive growth inwards at the subcutaneous inoculation site which might not be detected by callipers. Cell lines in the Control Dataset were categorised by metastatic profile. The pooled metastatic cell lines had smaller slope estimate values (less prominent morphology) at larger volumes in comparison to non-metastatic cell lines (Figure 4A). A Tukey’s Honest significant differences test on a fitted ANOVA concluded that there was no significant difference between the pooled metastatic and non-metastatic groups (p value = 0.5142). However, when metastatic and non-metastatic cell lines were split into volume bins and compared, results showed that there were significant differences in metastatic and non-metastatic tumours 100-200mm^3^ in size, and at volumes >1200mm^3^ (Table 4). Splitting results by cell line revealed different prominence trends within the metastatic and non-metastatic groups, indicating that metastatic potential was not the only variable affecting tumour prominence (Figure 4B).

**Figure 4.**
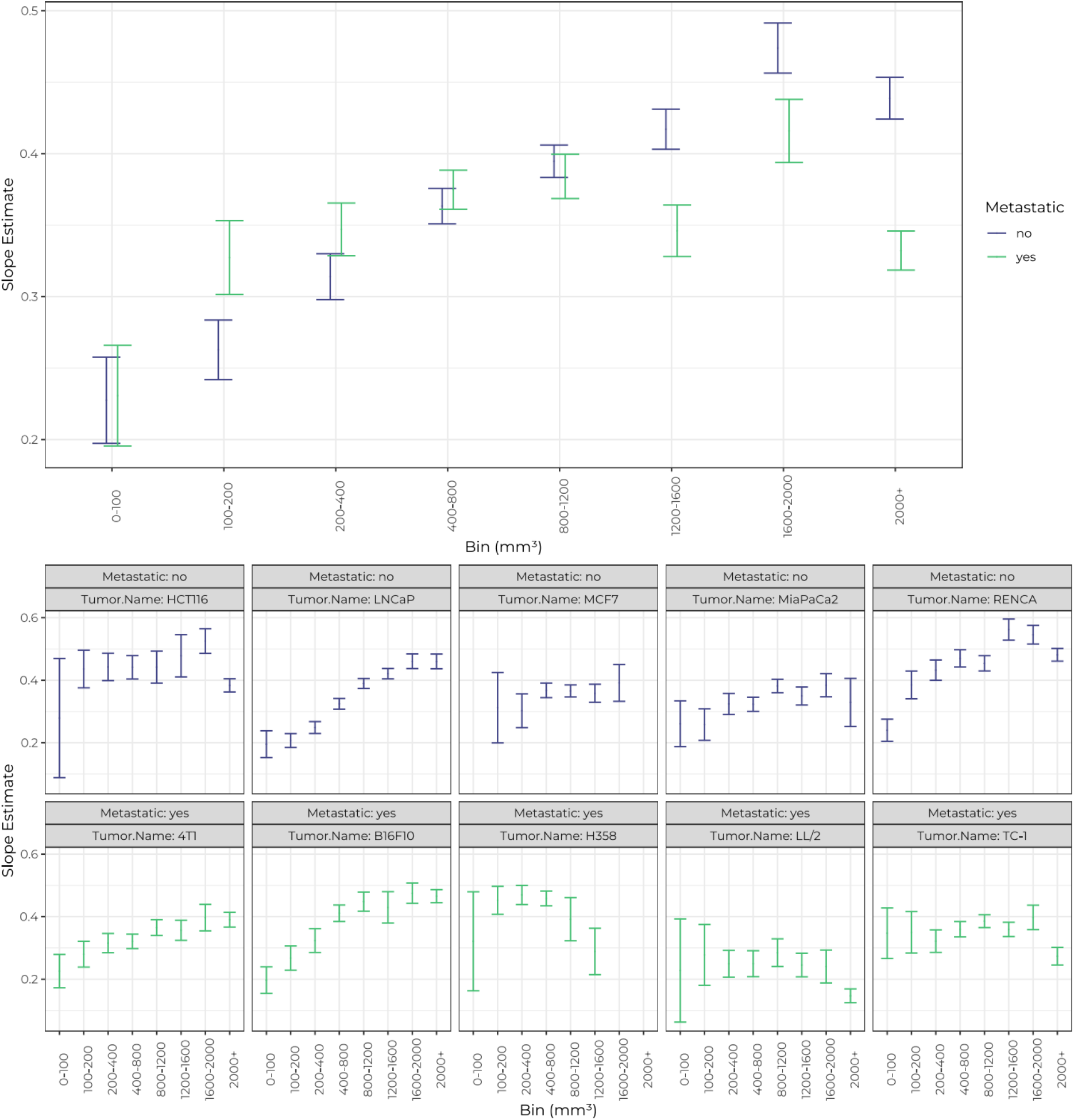
Slope estimates representing tumour prominence (Height divided by Width) for different tumour volumes. Error bars represent the 95% confidence intervals of the slope estimates. Metastatic (purple) and non-metastatic (green) groups and cell lines are indicated. **(A) Slope estimates for metastatic (n=1769) and non-metastatic (n=2380) tumours.** **(B) Individual slope estimates for 10 tumour cell lines.** Cell lines are labelled as metastatic or non-metastatic.

**Table 4.** Comparison of metastatic and non-metastatic tumour prominence across different volume ranges. A general linear hypothesis test was computed to test for significance between each of the metastatic and non-metastatic volume bins. P values rounded to 4 dp. Significance indicated: P>0.05 ns, P≤0.05 *, P≤0.005 **, P≤0.001 ***.

| Volume (mm<sup>3</sup>) | pvalue | Significance |
| --- | --- | --- |
| 0-100 | 1.0000 | n.s. |
| 100-200 | 0.0011 | ** |
| 200-400 | 0.0618 | n.s. |
| 400-800 | 0.8700 | n.s. |
| 800-1200 | 0.9263 | n.s. |
| 1200-1600 | < 0.0001 | *** |
| 1600-2000 | 0.0004 | *** |
| 2000+ | < 0.0001 | *** |

### Investigation of Tumour Type

Analysis of trends within and across tumour type groups was hindered by a lack of different cell lines of each tumour type, so carcinoma cell lines were compared against pooled non-carcinoma cell lines (Figure 5). The tumour prominence trends between these groups were significantly different overall (p < 0.0001, Table 5). Specifically, carcinomas differed in prominence from non-carcinoma tumours between 100-1600mm^3^. Tumours from the single lymphoma cell line (BL2) appeared flatter than average (data not shown), however more lymphoma cell lines need to be tested to confirm whether this trend is common to all lymphoma cell lines.

**Table 5.** Comparison of carcinoma and non-carcinoma tumour prominence across different volume ranges. A general linear hypothesis test was computed to test for significance between each of the carcinoma and non-carcinoma volume bins. P values rounded to 4 dp. Significance indicated: P>0.05 ns, P≤0.05 *, P≤0.005 **, P≤0.001 ***.

| Volume Bin (mm<sup>3</sup>) | P Value | Significance |
| --- | --- | --- |
| 0-100 | 0.3710 | ns |
| 100-200 | 0.0002 | *** |
| 200-400 | 0.0000 | *** |
| 400-800 | 0.0000 | *** |
| 800-1200 | 0.0036 | ** |
| 1200-1600 | 0.0003 | *** |
| 1600-2000 | 0.1100 | ns |
| 2000+ | 0.2840 | ns |

**Figure 5.**
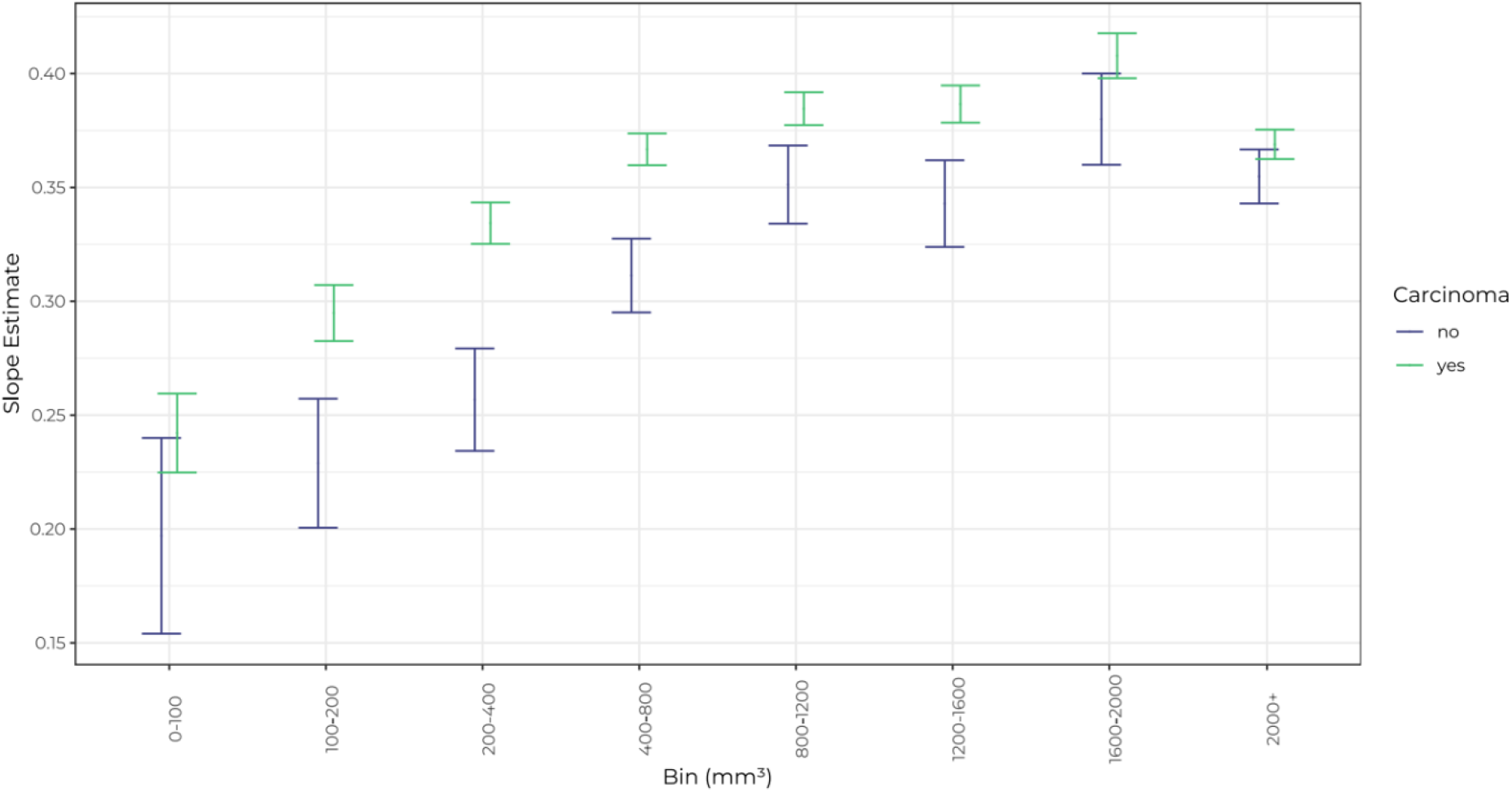
Slope estimates representing tumour prominence (Height divided by Width) for different carcinoma and non-carcinoma tumour volume ranges. Error bars represent the 95% confidence interval of the slope estimates. Number of scans: N=7652 (carcinomas) and N=1396 (non-carcinomas).

### Calliper vs 3D-TI Accuracy

Tumour prominence was compared to the accuracy of each measurement device. Excised tumour volumes were used to compare the accuracy of calliper measurement using the standard LWW formula, and 3D-TI using the LWH formula. The 3D-TI LWH volumes obtained a mean difference from excised tumour volume of −1.4% ± 4.2% whereas the LWW calliper volumes had a mean percentage difference of 50.5% ± 4.7% away from the excised weights, a difference 36X larger. 3D-TI volumes remained accurate at all levels of tumour prominence whereas the LWW calliper volumes became less accurate as prominence decreased. This was due to the overestimation of height by the LWW formula as previously shown.

### Simulating the Effect of Height Measurement on Efficacy Study Results

Simulated growth curves for LWW volumes were plotted so that the characteristics of 6 cell lines could be studied in more detail (Figure 7A). As expected, these simulated cell lines had different ‘growth’ rates, with RENCA growing at the quickest rate and LNCaP growing at the slowest rate. Rodent tumours were synthetically generated and the average synthetic rodent growth curves for LWW and LWH were plotted for each cell line (10 synthetic rodents per group, LNCaP example shown in Figure 7B). As expected, the LWH plots produced similar overall growth kinetics albeit at comparatively lower volumes than LWW.

**Figure 6.**
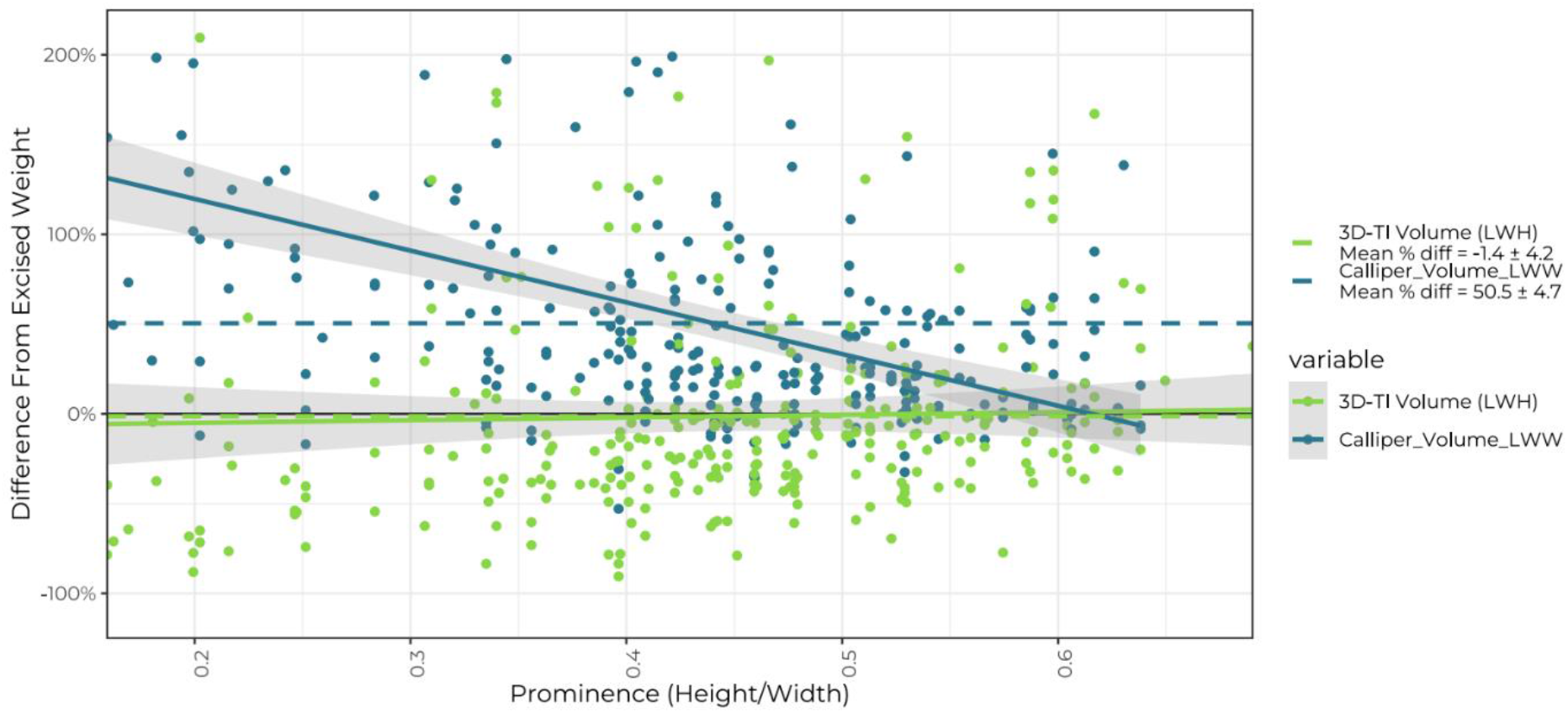
Accuracy of in vivo tumour measurement. Relative percentage accuracy was calculated by comparing in vivo tumour measurements to excised tumour volumes. In vivo volumes were calculated using 3D-TI LWH (green) and calliper LWW (blue) methods. Data points are ordered by prominence (Height/Width) values as recorded by 3D-TI. Trend lines (solid) were fitted using a linear model for each variable; shaded regions represent 95%CIs. Mean percentage difference for each method is shown as a dotted line (−1.4% ± 4.2% for 3D-TI LWH, 50.5% ± 4.7% for calliper LWW. Y axes cropped at −125% & 225%. N = 299 for 3D-TI, N = 274 for callipers.

**Figure 7.**
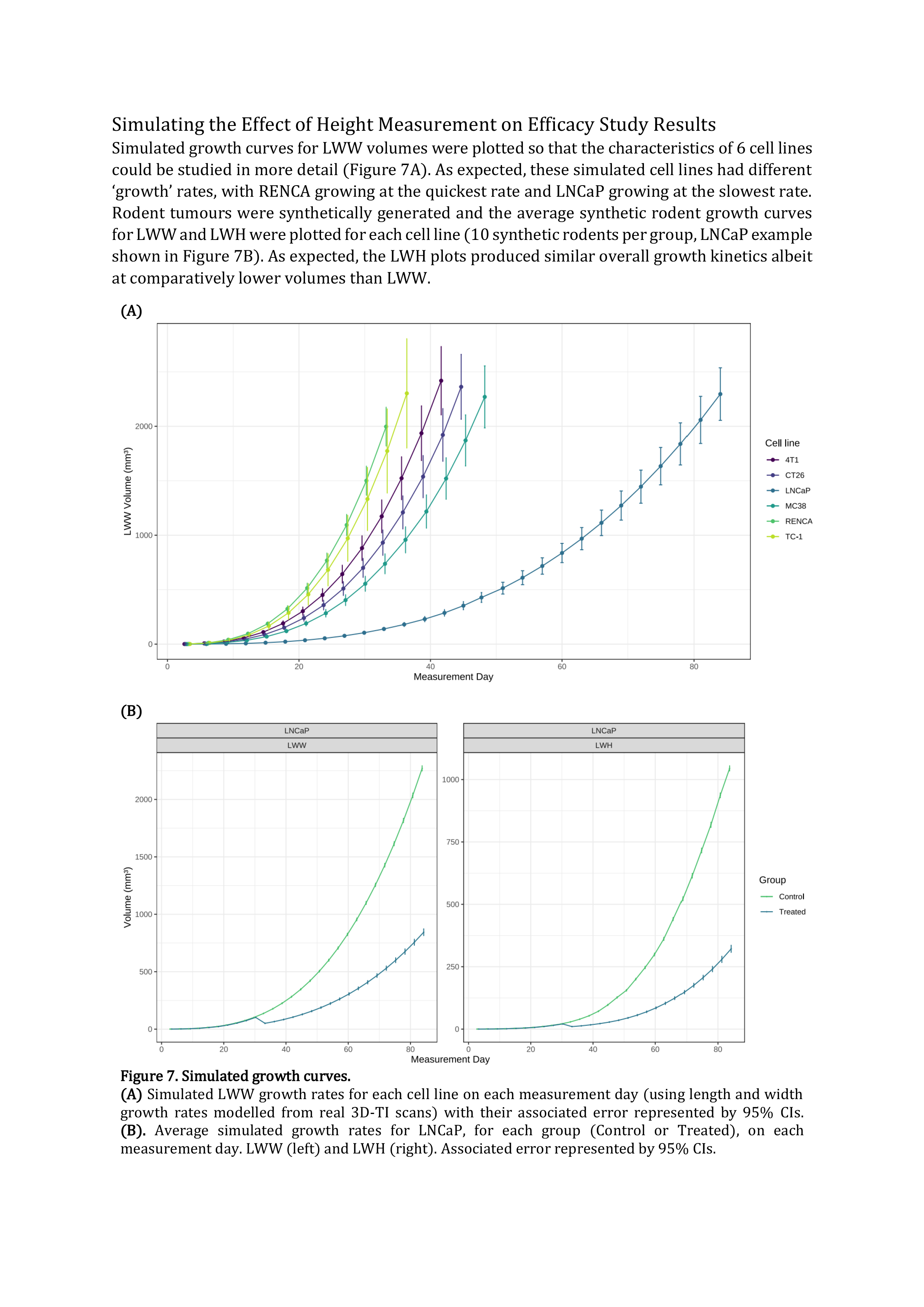
Simulated growth curves. **(A)**Simulated LWW growth rates for each cell line on each measurement day (using length and width growth rates modelled from real 3D-TI scans) with their associated error represented by 95% CIs. **(B).**Average simulated growth rates for LNCaP, for each group (Control or Treated), on each measurement day. LWW (left) and LWH (right). Associated error represented by 95% CIs.

TGI was calculated from the rodent ratios in the control and treated groups for each measurement method to determine treatment effect. The average TGI score from 250 repeats across 41 different treatment strengths was plotted for each of the cell lines and measurement methods (Figure 8A). TGI score was dependent on volume formula and cell line, as well as treatment strength. Therefore, different types of errors may be observed depending on the TGI threshold for effective treatment and the cell line concerned. For example, the LWH TGI curves for LNCaP, CT26, 4T1 and RENCA (for the most part) are lower than the LWW curve, meaning that TGI scores obtained using LWH will be smaller and therefore LWH could more easily detect a treatment effect in these cell lines. On the other hand, MC38 and TC-1 (greater than a certain treatment strength of ≈ 0.64) show the opposite effect, meaning that the use of LWW could incorrectly show a treatment effect when there is none. We conclude that the effect of including height in volume calculation on TGI is cell line-dependent.

**Figure 8.**
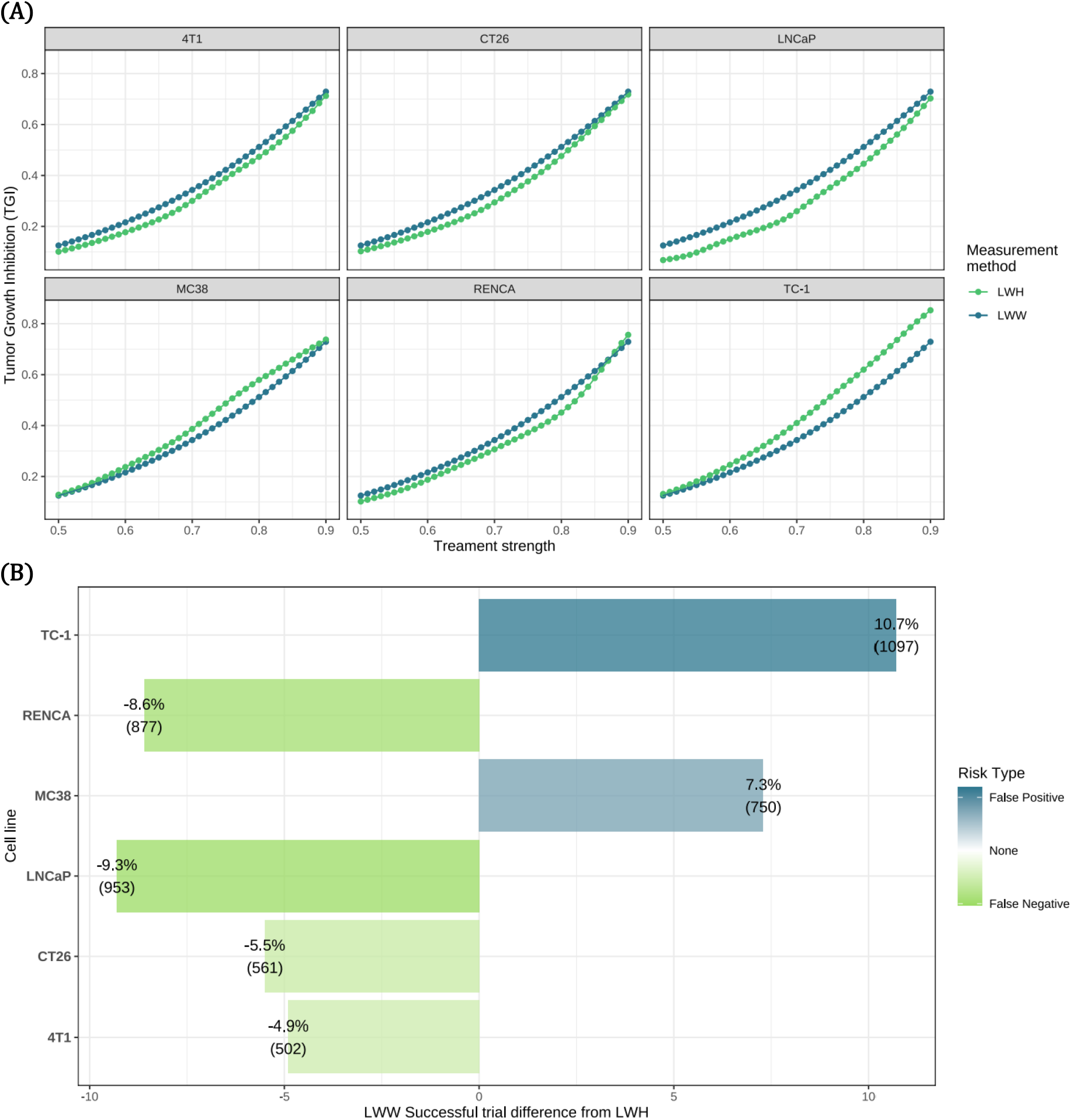
Simulated treatment effectiveness and error rates. **(A)** Average (mean) TGI values from 250 repeats per treatment level are plotted for LWW (blue) and LWH (green) measurement methods. 20 simulated animals were observed on each trial; 10 control and 10 treated. **(B)** The difference in number of successful trials for LWW compared to LWH (used as the gold standard) for each cell line, along with the associated risk type that these errors carry. Error types false positive (blue) and false negative (green) are shown.

These LWW errors came in the form of false positives or negatives dependent on the cell line and treatment level. A trial was considered a success when the TGI score was less than 0.45 and the relative percentage (and exact number) of false negatives and positives was determined for each cell line (Figure 8B). For LNCaP, more than 1 in 11 LWW simulated trials resulted in a false negative (9.3%) and for TC-1 more than 1 in 10 trials resulted in a false positive (10.7%) when compared to LWH. These results show that failure to measure tumour height can negatively affect conclusions from a trial.

## Discussion

Despite the current standard to use tumour width as a proxy for height in volume calculations, our analysis of 9,522 tumour scans showed that this is a poor approximation; tumour height is closer to one third of the tumour width on average. We found that tumour prominence changes throughout the growth cycle and furthermore this relationship is cell line-dependent, therefore methods using width as a proxy for height, or which approximate tumour height mathematically using a constant are not sufficient to accurately estimate tumour volume.

Comparison of 3D-TI LWH and calliper LWW volume measurements to excised tumour weights further confirmed that measurement using 3 dimensions is needed for greatest volumetric accuracy. The discrepancy between measurement methods increased as tumour prominence decreased, and we found that tumour prominence is lowest (tumours are flattest) at small volumes. This finding is particularly relevant for randomisation which typically occurs at tumour volumes between 75-150 mm^3^, a range that can be affected by the presence of hydrogels such as Matrigel, scarring, or inflammation at the site of tumour inoculation. Accuracy at small volumes is therefore crucial in order to confirm significant growth up until the point of randomisation, ensuring the mass being measured is indeed established tumour.

Determining the treatment window accurately at small volumes is also important during bioavailability and toxicity studies which require delivery of a therapeutic agent via the blood; the tumour must be large enough to be reliably established with a blood supply so that the drug can be delivered, but not too large that the optimum window for treatment has passed. Measuring height enables high accuracy and reproducibility when determining this point. 3D-TI not only provides more accurate results, but as previously shown, the automatic measurement feature reduces bias(15). Greater confidence at such small volumes is needed for correct set up of experiments from the very beginning, and to ensure data are reproducible.

6 carcinoma cell lines were investigated individually, and the relationship between prominence and tumour size varied for each. The 6 cell lines had high metastatic, low metastatic, or non-metastatic profiles (2 of each) but results did not reveal significant differences in prominence correlated to metastatic profile. This is likely due to the large variation of tumour morphology and growth kinetics found between different cell lines. Splitting the complete Control Dataset by tumour type revealed significant differences between carcinoma and non-carcinoma tumours however further analysis of a larger range of non-carcinoma cell lines is needed for more meaningful comparison. Two lymphoma cell lines did display below average prominence, however more lymphoma cell lines need to be studied to confirm if this effect is true for all subcutaneous lymphoma models.

Investigating a predicted relationship between invasive cell lines and tumour prominence further confirmed significant differences in prominence trends when individual cell lines within metastatic and non-metastatic groups were compared. Although best efforts were made to categorise cell lines’ metastatic potential based on available literature, reported experimental and data reporting methods varied, so it was difficult to definitively exclude cell lines as ‘not metastatic from subcutaneous sites’(37). We pooled data into metastatic and non-metastatic groups to provide a large sample size, and the results showed that overall, metastatic-type tumours were significantly less prominent (flatter) than non-metastatic tumours between volumes 100-200 mm^3^, and at volumes >1200mm^3^. We predict that at large volumes, this observation could be correlated to the switch to an invasive phenotype as tumour cells begin to invade below the basement membrane and spread to other tissues. We suggest that tracking subcutaneous tumour prominence could therefore be investigated as an early warning marker for metastasis; this would require further study as trends would likely be cell line specific.

Most tumour studies last for weeks and therefore follow tumours across a range of volumes as they grow and/or regress. As tumour prominence is dependent on cell line and tumour size, volumetric accuracy is affected if the LWW formula is used, but the effect is not uniform. For cell lines that produce subcutaneous tumours of constant prominence throughout the growth cycle like MC38 and LL/2, the LWW formula will consistently overestimate volumes. For others like TC-1, RENCA, and H358 where prominence increases then decreases as tumours grow, accuracy will increase, then decrease in correlation with tumour growth.

When using cell lines like LNCaP, B16F10 and CT26 that increase in prominence at larger volumes, accuracy when using LWW will be at its lowest at small volumes, increasing with tumour size. In particular, tumour models that follow this pattern are more likely to lead to more variable results in efficacy studies, as a decrease in measurement accuracy correlated with decreased size could mean that measurement of tumour regression is less accurate than measurement of tumour growth. Lower accuracy at smaller volumes also impacts randomisation and dosing start points as previously discussed. These variations in accuracy contribute to experimental variation and irreproducibility of data. By using modelling to look at the effects of measurement inaccuracy caused by using the LWW formula vs LWH, we found that the failure to measure tumour height can result in more false positive or negative results. False positive results represent ineffective drugs being brought to clinical trial, wasting time and money in the process. Conversely, false negative results represent missing a successful treatment effect and losing an effective drug candidate.

### Future Considerations

Callipers remain the most commonly used tool for tumour measurement but are rarely used to measure tumour height due to difficulty of the technique causing measurement variability even higher than when measuring length and width(2,38). 3D-TI can be used to measure tumour height quickly and accurately, without reagents or anaesthesia so is an attractive alternative to current measurement devices to achieve accurate tumour volumes. Because all tumour measurements are captured from a single scan, measuring 3 dimensions is no slower than measuring 2, in contrast to measurements with callipers, where every additional measurement increases the measurement session and rodent handling time.

Because 3D-TI reconstructs whole tumours digitally as a 3D model, even greater accuracy could be achieved by calculating true volume and area. Although the 3D-TI, MRI, ultrasound, and other imaging technologies are ready for these innovations, the cancer research community is wary of moving too far away from the LWW formula upon which current welfare guidelines, dosing schemes, and randomisation points are based. Conversion models exist to solve these issues, however more support at institutional levels, time, and funding are needed to implement new technologies and methods.

We have shown that many tumour cell lines vary in prominence as subcutaneous tumour volume changes, and that height is a unique parameter independent of tumour width. For that reason, we strongly advise against using the LWW formula, or other formulae that exclude tumour height. In our opinion, the best practice for accurate tumour volume measurements is by including height, using the LWH formula. Due to the reproducibility crisis affecting oncology drug trials, we cannot afford to lose data by ignoring tumour height when there are methods available to measure it.

### Contribution to the field

Our work, taken from a large dataset of real-world tumour measurements highlights a flaw in the most commonly used methodology for monitoring subcutaneous tumour volumes. The assumption that tumour width is equal to height is incorrect, resulting in overestimation of tumour volume. Changes in tumour height are not recorded using the LWW volume calculation, and we have shown that height and width are not linearly correlated, so by ignoring height, information about treatment effect is lost. We also showed that 3D-TI measurement of height, length and width achieves greater accuracy when compared to a common calliper method.3D-TI also has advantages in comparison to existing imaging methods like ultrasound and MRI that offer improved accuracy in comparison to callipers including higher throughput, faster acquisition, and no reliance on reagents or anaesthesia.

Measuring treatment efficacy by monitoring subcutaneous tumour growth in vivo is a key step in preclinical studies. The oncology field has the lowest rate of drug development success, due to poor reproducibility and translation. More accurate results in preclinical stages will increase confidence in drug candidates chosen to progress to clinical trials.

Tumour height is an independent variable that is largely ignored by scientists performing in vivo trials using callipers. Due to the reproducibility crisis affecting oncology drug trials, we cannot afford to lose data by ignoring tumour height when there are methods available to measure it.

## Acknowledgements

We thank Elisa Heyrman for her advice and edits to the manuscript.

## Abbreviations List

3D-TI: 3D and thermal imaging
ANOVA: Analysis of variance
CT: Computerised tomography
GAM: Generalised additive model
LWH: Volume formula using length, width, and height
LWW: Volume formula using length and width
MRI: Magnetic resonance imaging
TGI: Tumour growth inhibition

## Bibliography

1. Euhus DM, Hudd C, Laregina MC, Johnson FE. Tumor measurement in the nude mouse. Journal of Surgical Oncology. 1986;31(4).

2. Dethlefsen LA, Prewitt JMS, Mendelsohn ML. Analysis of tumor growth curves. Journal of the National Cancer Institute. 1968;40(2).

3. Sápi J, Kovács L, Drexler DA, Kocsis P, Gajári D, Sápi Z. Tumor Volume Estimation and Quasi-Continuous Administration for Most Effective Bevacizumab Therapy. PLoS One. 2015;10(11):e0142190.

4. Lee JC, Lee H, Moon C, Lee S, Lee W, Cha H, et al. Arsenic Trioxide as a Vascular Disrupting Agent: Synergistic Effect with Irinotecan on Tumor Growth Delay in a CT26 Allograft Model. Translational oncology. 2013 Feb 1;6:83–91.

5. Nguyen DX, Bos PD, Massagué J. Metastasis: from dissemination to organ-specific colonization. Nat Rev Cancer. 2009 Apr;9(4):274–84.

6. Fontaine JJ, Marangoni E, Chateau-Joubert S, Servely JL. Pathology of Patient-Derived Xenograft Tumors. In: Patient Derived Tumor Xenograft Models [Internet]. Elsevier; 2017 [cited 2022 Jun 28]. p. 135–48. Available from: https://linkinghub.elsevier.com/retrieve/pii/B9780128040102000102

7. Jensen MM, Jørgensen JT, Binderup T, Kjær A. Tumor volume in subcutaneous mouse xenografts measured by microCT is more accurate and reproducible than determined by 18F-FDG-microPET or external caliper. BMC Medical Imaging. 2008;8.

8. Tomayko MM, Reynolds CP. Determination of subcutaneous tumor size in athymic (nude) mice. Cancer Chemotherapy and Pharmacology. 1989;24(3).

9. Feldman JP, Goldwasser R. a Mathematical Model for Tumor Volume Evaluation Using Two-Dimensions. Jpurnal of applied quantitative methods. 2009;4(4).

10. Hall C, Grabowiecki Y von, Pearce SP, Dive C, Bagley S, Muller PAJ. iRFP (near-infrared fluorescent protein) imaging of subcutaneous and deep tissue tumours in mice highlights differences between imaging platforms. Cancer Cell International. 2021;21(1).

11. Ni J, Bongers A, Chamoli U, Bucci J, Graham P, Li Y. In Vivo 3D MRI Measurement of Tumour Volume in an Orthotopic Mouse Model of Prostate Cancer. Cancer Control. 2019;26(1).

12. Ayers GD, McKinley ET, Zhao P, Fritz JM, Metry RE, Deal BC, et al. Volume of preclinical xenograft tumors is more accurately assessed by ultrasound imaging than manual caliper measurements. Journal of Ultrasound in Medicine. 2010;29(6).

13. Delgado San Martin J, Ehrhardt B, Paczkowski M, Hackett S, Smith A, Waraich W, et al. An innovative non-invasive technique for subcutaneous tumour measurements. PLoS ONE. 2019;14(10).

14. Murkin JT, Amos HE, Brough DW, Turley KD. In silico modelling demonstrates that user variability during tumor measurement can affect in vivo therapeutic efficacy outcomes [Internet]. bioRxiv; 2022 [cited 2022 Aug 22]. p. 2022.04.20.487864. Available from: https://www.biorxiv.org/content/10.1101/2022.04.20.487864v1

15. Brough D, Smith A, Turley K, Amos H, Murkin J. Reducing inter-operator variability when measuring subcutaneous tumours in mice: An investigation into the effect of inter-operator variability when using callipers or a novel 3D and thermal measurement system. MetaArXiv Preprints. 2021 Oct 8;

16. Thiel DD, Jorns J, Lohse CM, Cheville JC, Thompson RH, Parker AS. Maximum tumor diameter is not an accurate predictor of renal cell carcinoma tumor volume. Scandinavian Journal of Urology. 2013;47(6).

17. Workman P, Aboagye EO, Balkwill F, Balmain A, Bruder G, Chaplin DJ, et al. Guidelines for the welfare and use of animals in cancer research. Br J Cancer. 2010 May;102(11):1555–77.

18. mgcv: Mixed GAM Computation Vehicle with Automatic Smoothness Estimation version 1.8-40 from CRAN [Internet]. [cited 2022 Jul 18]. Available from: https://rdrr.io/cran/mgcv/

19. Wickham H. ggplot2: Elegant Graphics for Data Analysis [Internet]. Springer-Verlag New York; 2016. Available from: https://ggplot2.tidyverse.org

20. Wood SN. Fast stable restricted maximum likelihood and marginal likelihood estimation of semiparametric generalized linear models. Journal of the Royal Statistical Society (B). 2011;73(1):3–36.

21. R Core Team. R: A Language and Environment for Statistical Computing [Internet]. Vienna, Austria: R Foundation for Statistical Computing; 2021. Available from: https://www.R-project.org/

22. Huang L, Li Y, Du Y, Zhang Y, Wang X, Ding Y, et al. Mild photothermal therapy potentiates anti-PD-L1 treatment for immunologically cold tumors via an all-in-one and all-in-control strategy. Nat Commun. 2019 Oct 25;10(1):4871.

23. Fox J, Weisberg S. An R Companion to Applied Regression [Internet]. Third. Thousand Oaks CA: Sage; 2019. Available from: https://socialsciences.mcmaster.ca/jfox/Books/Companion/

24. Masuda J, Takayama E, Strober W, Satoh A, Morimoto Y, Honjo Y, et al. Tumor growth limited to subcutaneous site vs tumor growth in pulmonary site exhibit differential effects on systemic immunities. Oncology Reports. 2017 Jan;38(1):449–55.

25. Terracina KP, Aoyagi T, Huang WC, Nagahashi M, Yamada A, Aoki K, et al. Development of a metastatic murine colon cancer model. Journal of Surgical Research. 2015 Nov;199(1):106–14.

26. Corbett TH, Griswold DPJr, Roberts BJ, Peckham JC, Schabel FM Jr. Tumor Induction Relationships in Development of Transplantable Cancers of the Colon in Mice for Chemotherapy Assays, with a Note on Carcinogen Structure1. Cancer Research. 1975 Sep 1;35(9):2434–9.

27. Gao ZG, Tian L, Hu J, Park IS, Bae YH. Prevention of metastasis in a 4T1 murine breast cancer model by doxorubicin carried by folate conjugated pH sensitive polymeric micelles. Journal of Controlled Release. 2011 May;152(1):84–9.

28. Yang S, Zhang JJ, Huang XY. Mouse Models for Tumor Metastasis. In: Rational Drug Design: Methods and Protocols [Internet]. Totowa, NJ: Humana Press; 2012 [cited 2022 Sep 26]. p. 221–8. (Methods in Molecular Biology). Available from: https://doi.org/10.1007/978-1-62703-008-3_17

29. Wu L, Wang W, Dai M, Li H, Chen C, Wang D. PPARα ligand, AVE8134, and cyclooxygenase inhibitor therapy synergistically suppress lung cancer growth and metastasis. BMC Cancer. 2019 Dec 2;19(1):1166.

30. Thalmann GN, Anezinis PE, Chang SM, Zhau HE, Kim EE, Hopwood VL, et al. Androgen-independent cancer progression and bone metastasis in the LNCaP model of human prostate cancer. Cancer Res. 1994 May 15;54(10):2577–81.

31. Murphy KA, James BR, Wilber A, Griffith TS. A Syngeneic Mouse Model of Metastatic Renal Cell Carcinoma for Quantitative and Longitudinal Assessment of Preclinical Therapies. J Vis Exp. 2017 Apr 12;(122).

32. Hothorn T, Bretz F, Westfall P. Simultaneous Inference in General Parametric Models. Biometrical Journal. 2008;50(3):346–63.

33. Gao J, Li N, Dong Y, Li S, Xu L, Li X, et al. miR-34a-5p suppresses colorectal cancer metastasis and predicts recurrence in patients with stage II/III colorectal cancer. Oncogene. 2015 Jul;34(31):4142–52.

34. Nagaraja GM, Othman M, Fox BP, Alsaber R, Pellegrino CM, Zeng Y, et al. Gene expression signatures and biomarkers of noninvasive and invasive breast cancer cells: comprehensive profiles by representational difference analysis, microarrays and proteomics. Oncogene. 2006 Apr;25(16):2328–38.

35. Gradiz R, Silva HC, Carvalho L, Botelho MF, Mota-Pinto A. MIA PaCa-2 and PANC-1 – pancreas ductal adenocarcinoma cell lines with neuroendocrine differentiation and somatostatin receptors. Sci Rep. 2016 Feb 17;6(1):1–14.

36. Basu I, Locker J, Cassera MB, Belbin TJ, Merino EF, Dong X, et al. Growth and Metastases of Human Lung Cancer Are Inhibited in Mouse Xenografts by a Transition State Analogue of 5’-Methylthioadenosine Phosphorylase. J Biol Chem. 2011 Feb 11;286(6):4902–11.

37. Fitzgerald JE, Byrd BK, Patil RA, Strawbridge RR, Davis SC, Bellini C, et al. Heterogeneity of circulating tumor cell dissemination and lung metastases in a subcutaneous Lewis lung carcinoma model. Biomed Opt Express. 2020 Jun 8;11(7):3633–47.

38. Thalheimer RD, Merker VL, Ly KI, Champlain A, Sawaya J, Askenazi NL, et al. Validating Techniques for Measurement of Cutaneous Neurofibromas: Recommendations for Clinical Trials. Neurology. 2021 Aug 17;97(7 Supplement 1):S32–41.

