## Supplementary Information for "Trends in Subcutaneous Tumour Height and Impact on Measurement Accuracy"

#### Supplementary Tables

| Cell Line | Tumour Type | Metastatic | Scans |
|---|---|---|---|
| **HCT116** | Carcinoma | no | 188 |
| **LNCaP** | Carcinoma | no | 985 |
| **MCF7** | Carcinoma | no | 228 |
| **MiaPaCa2** | Carcinoma | no | 376 |
| **RENCA** | Carcinoma | no | 607 |
| **4T1** | Carcinoma | yes | 481 |
| **B16F10** | Melanoma | yes | 465 |
| **H358** | Carcinoma | yes | 243 |
| **LL/2** | Carcinoma | yes | 176 |
| **TC-1** | Carcinoma | yes | 419 |

**Table S1.** Cell lines included in the pooled metastatic analysis.

| Tumour Type | Tumour Name | Scans |
|---|---|---|
| **Carcinoma** | 4T1 | 481 |
| **Carcinoma** | 786-O | 18 |
| **Melanoma** | A375 | 36 |
| **Carcinoma** | A431 | 20 |
| **Carcinoma** | A549 | 360 |
| **Carcinoma** | AR42J | 453 |
| **Melanoma** | B16 | 76 |
| **Melanoma** | B16F10 | 465 |
| **Lymphoma** | BL2 | 380 |
| **Carcinoma** | BR5FVB1 | 10 |
| **Carcinoma** | CT26 | 657 |
| **Carcinoma** | H1299 | 267 |
| **Carcinoma** | H1975 | 47 |
| **Carcinoma** | H358 | 243 |
| **Carcinoma** | HCC1806 | 185 |
| **Carcinoma** | HCC1954 | 10 |
| **Carcinoma** | HCT116 | 188 |
| **Leukaemia** | HL-60 | 107 |
| **Carcinoma** | HT-29 | 90 |
| **Carcinoma** | KP1.9 | 66 |
| **Carcinoma** | LL/2 | 176 |
| **Carcinoma** | LNCaP | 985 |
| **Carcinoma** | MC38 | 1020 |
| **Carcinoma** | MCF7 | 228 |
| **Melanoma** | MDA-MB-453 | 10 |

| | | |
|---|---|---|
| **Leukaemia** | MOLM-13 | 15 |
| **Carcinoma** | MiaPaCa2 | 376 |
| **Carcinoma** | NCI-H1299 | 354 |
| **Carcinoma** | NCI-H2073 | 137 |
| **Carcinoma** | NCI-H522 | 50 |
| **Carcinoma** | NCI-H82 | 130 |
| **Leukaemia** | OCI-AML3 | 86 |
| **Carcinoma** | Pa-Tu-8902 | 75 |
| **Carcinoma** | RENCA | 607 |
| **Carcinoma** | TC-1 | 419 |
| **Lymphoma** | TMD8 | 12 |
| **Glioma** | U87 | 209 |

**Table S2.** Cell lines and scans included in the analysis by tumour type (carcinoma vs other).

| Combination | Difference | Lower | Upper | pvalue | Significance |
|---|---|---|---|---|---|
| **CT26-4T1** | 0.050401 | 0.030453 | 0.07035 | < 0.0001 | *** |
| **LNCaP-4T1** | -0.01164 | -0.0301 | 0.006823 | 0.4670 | ns |
| **MC38-4T1** | 0.013075 | -0.00538 | 0.031527 | 0.3310 | ns |
| **RENCA-4T1** | 0.075032 | 0.054771 | 0.095293 | < 0.0001 | *** |
| **TC-1-4T1** | 0.054604 | 0.03197 | 0.077238 | < 0.0001 | *** |
| **LNCaP-CT26** | -0.06204 | -0.07874 | -0.04534 | < 0.0001 | *** |
| **MC38-CT26** | -0.03733 | -0.05402 | -0.02063 | < 0.0001 | *** |
| **RENCA-CT26** | 0.024631 | 0.005958 | 0.043304 | 0.0024 | ** |
| **TC-1-CT26** | 0.004202 | -0.01702 | 0.025427 | 0.9930 | ns |
| **MC38-LNCaP** | 0.024714 | 0.009829 | 0.039599 | < 0.0001 | *** |
| **RENCA-LNCaP** | 0.086671 | 0.069596 | 0.103747 | < 0.0001 | *** |
| **TC-1-LNCaP** | 0.066243 | 0.046409 | 0.086076 | < 0.0001 | *** |
| **RENCA-MC38** | 0.061957 | 0.044892 | 0.079023 | < 0.0001 | *** |
| **TC-1-MC38** | 0.041529 | 0.021704 | 0.061354 | < 0.0001 | *** |
| **TC-1-RENCA** | -0.02043 | -0.04195 | 0.00109 | 0.0741 | ns |

**Table S3.** A Tukey honest significant differences test was computed using the fitted ANOVA to give a multiple pairwise-comparison between the group means. P values rounded to 4 dp. Significance indicated: P>0.05 ns, P≤0.05 *, P≤0.005 **, P≤0.001 ***

### Supplementary Figures: Model Validation

**(A)**

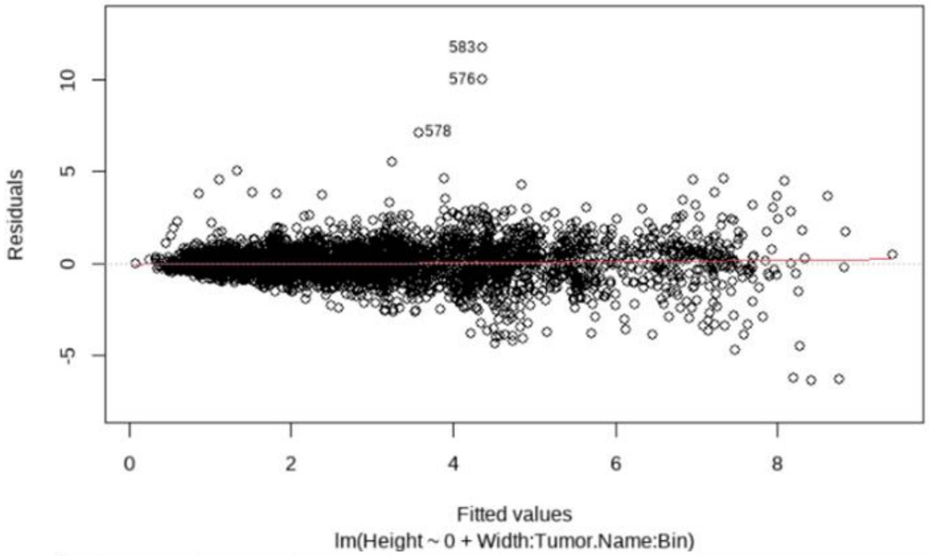


**(B)**

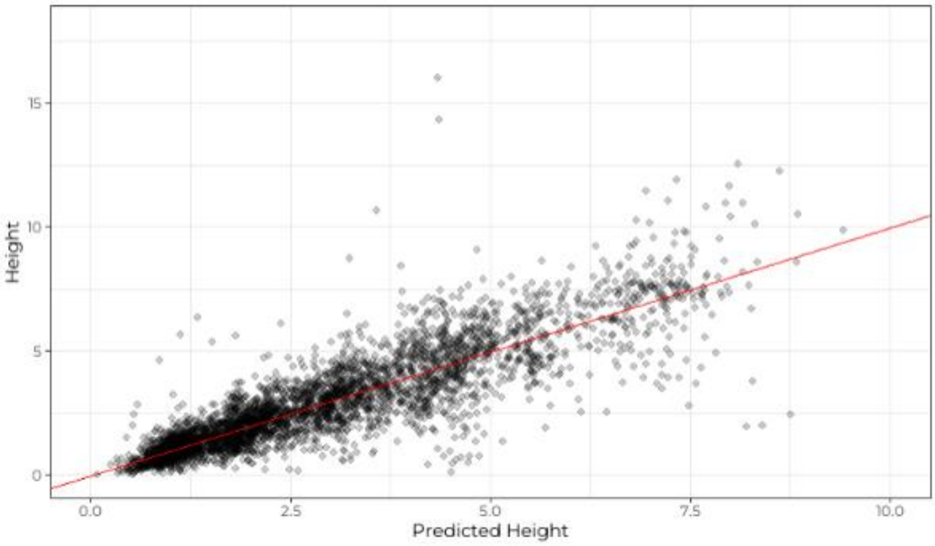


**Figure S1A**. A scatter plot of the residuals against their fitted values for the Height=Width:TumourName:Bin model formula for cell lines 4T1, CT26, LNCaP, MC38, RENCA, and TC-1. The red line indicates the mean residual value for every fitted value region. Linearity holds when the red line is close to the dashed (grey) line. If homoskedacity holds, the spread of the residuals is the constant across the x-axis. Outliers are also labelled.

**S1B.** Actual vs predicted values from the Height = Width:TumourName:Bin model formula. In red is a reference y = x line.

(A)

(B)

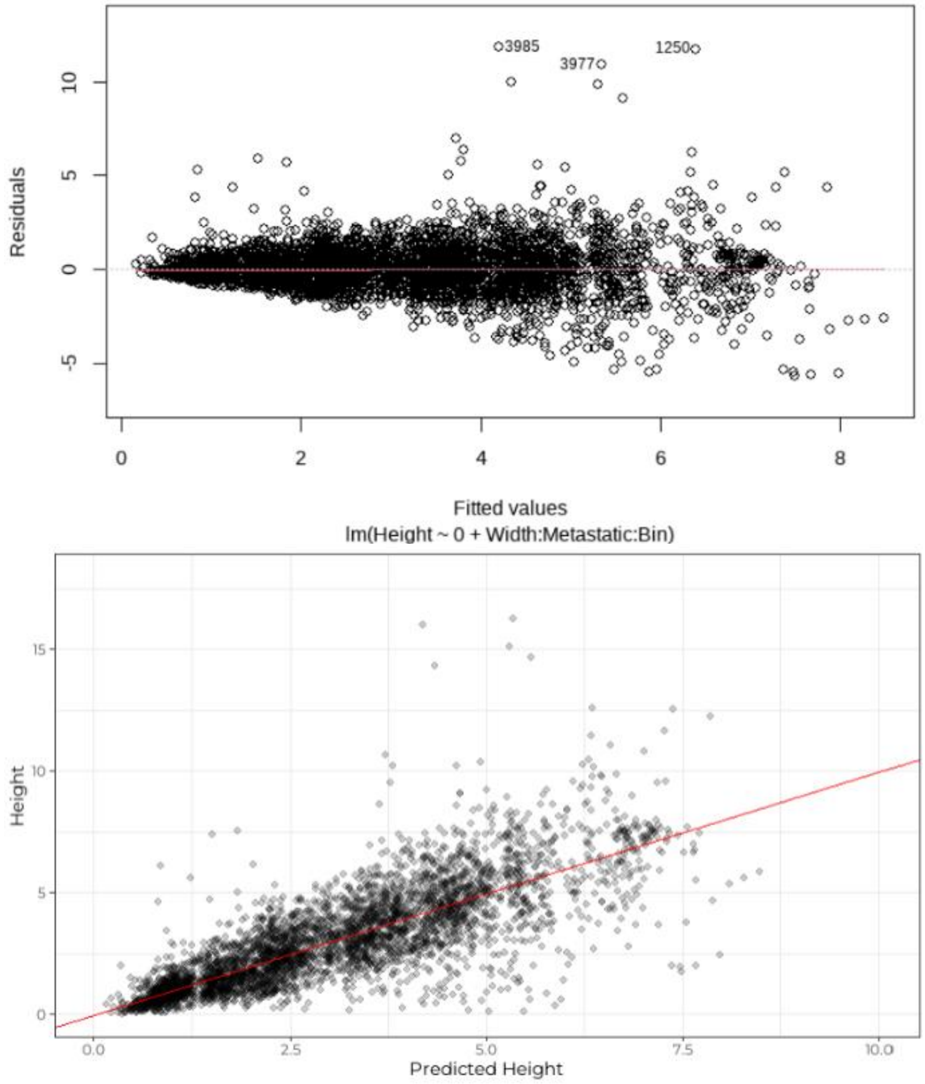


**Figure S2A.** A scatter plot of the residuals against their fitted values for the Height = Width:Metastatic:Bin model formula. The red line indicates the mean residual value for every fitted value region. Linearity holds when the red line is close to the dashed (grey) line. If homoskedacity holds, the spread of the residuals is the constant across the x-axis. Outliers are also labelled.
**S2B.** Actual vs predicted values from the Height = Width:Metastatic:Bin model formula. In red is a reference y = x line.

(A)

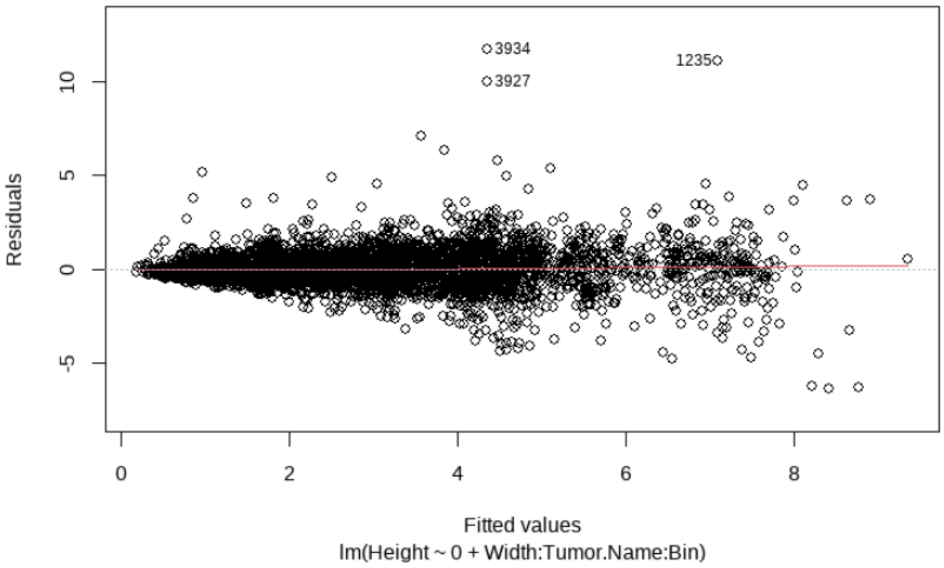


(B)

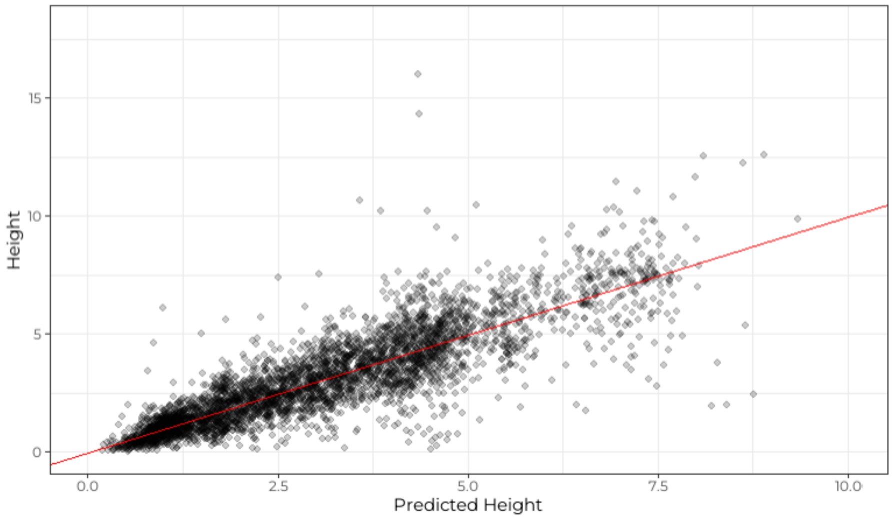


**Figure S3A.** A scatter plot of the residuals against their fitted values for the Height = Width:Tumour.Name:Bin model formula for 5 metastatic and 5 non-metastatic cell lines. The red line indicates the mean residual value for every fitted value region. Linearity holds when the red line is close to the dashed (grey) line. If homoskedacity holds, the spread of the residuals is the constant across the x-axis. Outliers are also labelled.

**S3B.** Actual vs predicted values from the Height = Width:TumourName:Bin model formula. In red is a reference y = x line.

(A)

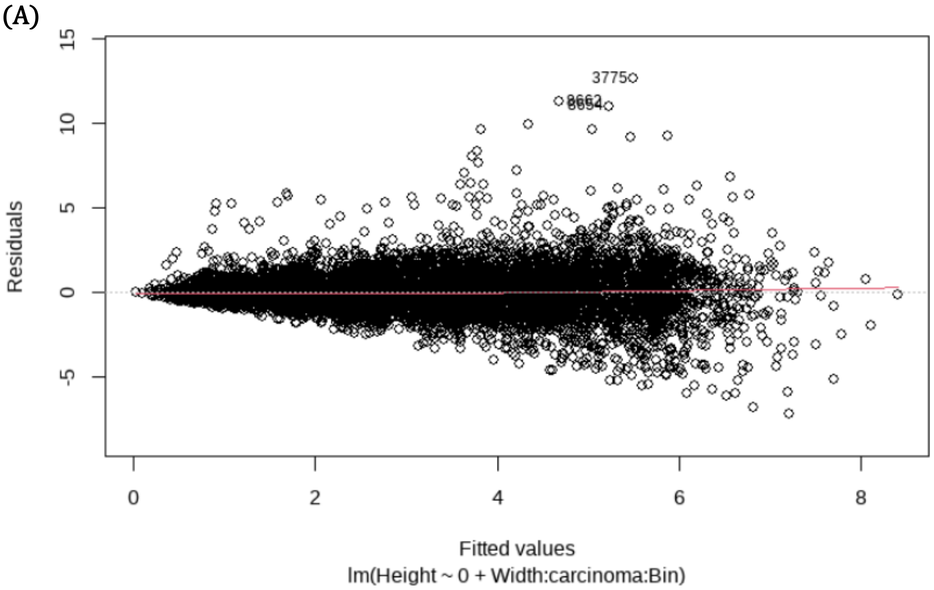


(B)

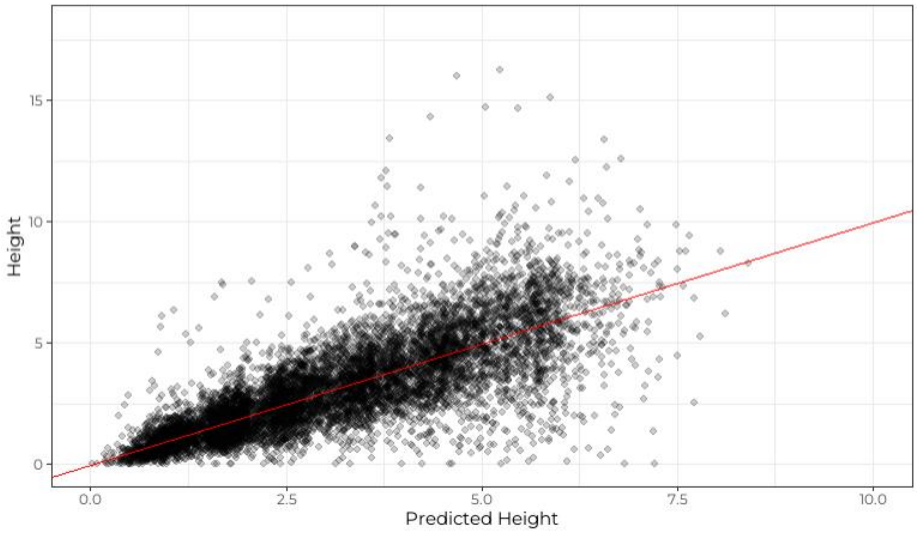


**Figure S4A.** A scatter plot of the residuals against their fitted values for the Height = Width:carcinoma:Bin model formula. The red line indicates the mean residual value for every fitted value region. Linearity holds when the red line is close to the dashed (grey) line. If homoskedacity holds, the spread of the residuals is the constant across the x-axis. Outliers are also labelled.

**B.** Actual vs predicted values from the Height = Width:carcinoma:Bin model formula. In red is a reference y = x line.

(A)

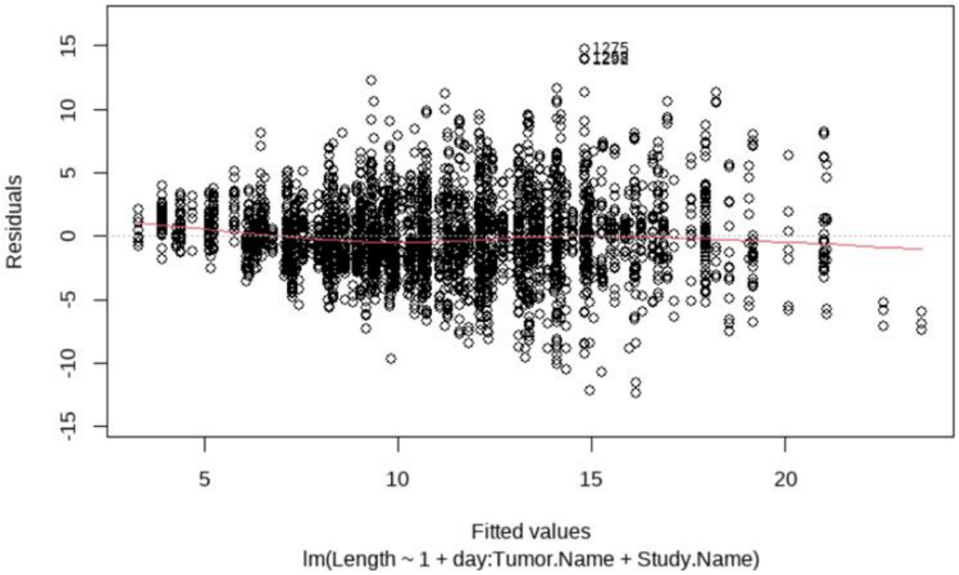


(B)

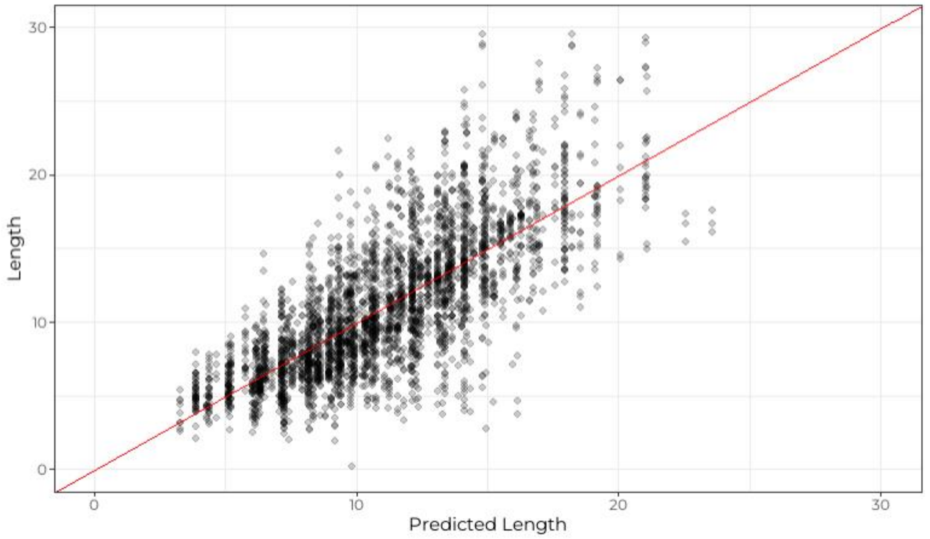


**Figure S5A.** A scatter plot of the residuals against their fitted values for the Length = day:Tumour.Name + Study.Name model formula. The red line indicates the mean residual value for every fitted value region. Linearity holds when the red line is close to the dashed (grey) line. If homoskedacity holds, the spread of the residuals is the constant across the x-axis. Outliers are also labelled.

**B,** Actual vs predicted values from the Length = day:Tumour.Name + Study.Name model formula. In red is a reference y = x line.

(A)

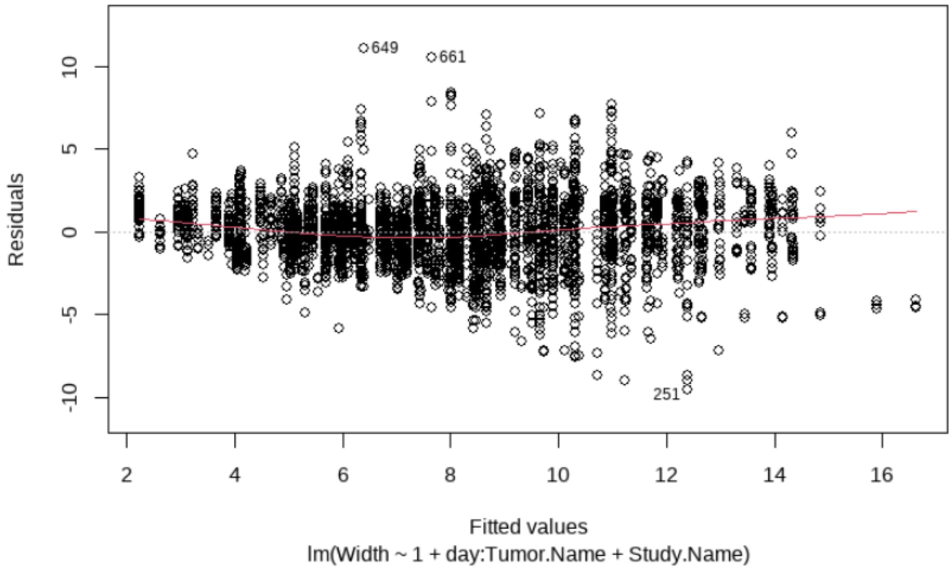


(B)

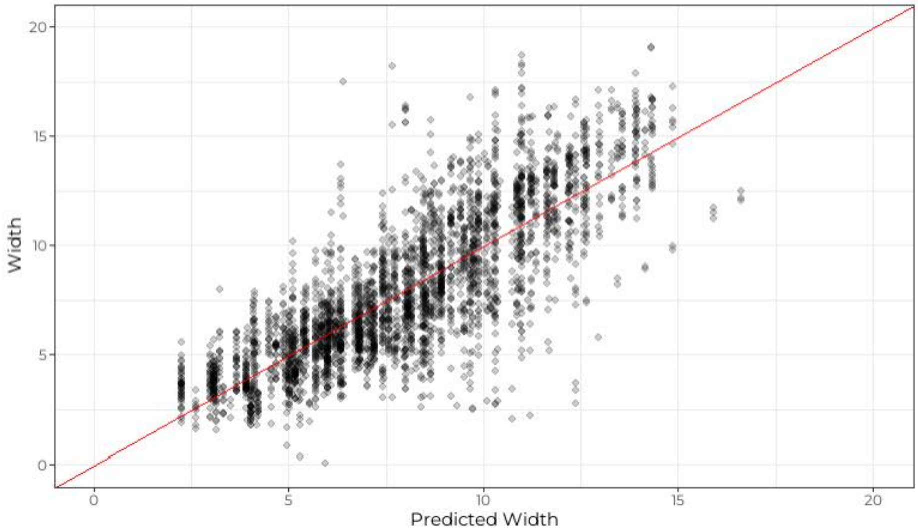


**Figure S6A.** A scatter plot of the residuals against their fitted values for the **Width = day:Tumour.Name + Study.Name** model formula. The red line indicates the mean residual value for every fitted value region. Linearity holds when the red line is close to the dashed (grey) line. If homoskedacity holds, the spread of the residuals is the constant across the x-axis. Outliers are also labelled.

**B.** Actual vs predicted values from the **Width = day:Tumour.Name + Study.Name** model formula. In red is a reference y = x line.
